# Late life metformin treatment limits cell survival and shortens lifespan by triggering an aging-associated failure of energy metabolism

**DOI:** 10.1101/863357

**Authors:** Lilia Espada, Alexander Dakhovnik, Prerana Chaudhari, Asya Martirosyan, Laura Miek, Tetiana Poliezhaieva, Yvonne Schaub, Ashish Nair, Nadia Döring, Norman Rahnis, Oliver Werz, Andreas Koeberle, Joanna Kirkpatrick, Alessandro Ori, Maria A. Ermolaeva

## Abstract

The diabetes drug metformin is to be clinically tested in aged humans to achieve health span extension, but little is known about responses of old non-diabetic individuals to this drug. By *in vitro* and *in vivo* tests we found that metformin shortens life span and limits cell survival when provided in late life, contrary to its positive early life effects. Mechanistically, metformin exacerbates aging-associated mitochondrial dysfunction towards respiratory failure, aggravated by the inability of old cells to upregulate glycolysis in response to metformin, leading to ATP exhaustion. The beneficial dietary restriction effect of metformin on lipid reserves is abrogated in old animals, contributing to metabolic failure, while ectopic stabilization of cellular ATP levels alleviates late life metformin toxicity *in vitro* and *in vivo*. The toxicity is also suspended in nematodes carrying diabetes-like insulin receptor insufficiency and showing prolonged resilience to metabolic stress induced by metformin. In sum, we uncovered an alarming metabolic decay triggered by metformin in late life which may limit its benefits for non-diabetic elderly patients. Novel regulators of life extension by metformin are also presented.

**Highlights:**

- Late life metformin treatment limits cell survival and shortens lifespan.
- Metformin exacerbates aging-associated mitochondrial dysfunction causing fatal ATP exhaustion.
- Old cells fail to upregulate glycolysis as a compensatory response to metformin.
- The dietary restriction (DR) mimetic response to metformin is abrogated in old animals.
- PKA and not AMPK pathway instigates the early life DR response to metformin.
- Stabilization of cellular ATP levels alleviates late life metformin toxicity *in vitro* and *in vivo*.

## Introduction

Metformin is among the most frequently prescribed drugs worldwide and it is used to facilitate glucose catabolism in patients with impaired insulin signaling (diabetes type 2) (Salani et al., 2014). Metformin is thought to act by inhibiting mitochondrial respiration (Wheaton et al., 2014). Recently, metformin was tested for additional physiological effects and found to extend life span in animal models ranging from nematodes to mice (Martin-Montalvo et al., 2013; Onken and Driscoll, 2010). Extensive clinical use in humans enabled the collection and analysis of data on the longevity of human diabetes patients treated with metformin. The metformin-exposed diabetes cohort was found to be longer lived than untreated healthy subjects (Bannister et al., 2014), in line with potential life-prolonging properties of metformin.

Type 2 diabetes is an aging-associated disorder and many patients start metformin treatment in late life. Based on survival analysis of diabetes patients, it was proposed that life prolonging effects of metformin may extend also to metabolic-healthy elderly individuals. Considering moderate metformin side effects in diabetes and the potential healthy aging benefits, metformin has emerged as an attractive candidate to be clinically tested as the first prospective anti-aging drug in humans. A short term trial administering metformin to aged (≥65 years old) pre-diabetic humans for a period of 6 weeks had recently been completed and the data was reported (Kulkarni et al., 2018). While effects on pathways such as TOR and immune response were observed, no obvious physiological changes were detected due to short duration of the treatment, leaving long-term effects of metformin on aged healthy humans an open question. This question is however critical because non-diabetic elderly humans are anticipated to be the first recipients of the putative health span extension treatment with metformin.

Via literature research of animal studies providing evidence of longevity modulation by metformin, we discovered that the pro-survival effect of this drug was mostly studied in young animals or animals exposed to metformin from young adulthood. The few studies performed in older animals (56-60 week old mice, average lifespan 96 weeks; 8 day old nematodes, average lifespan 14-21 days), either failed to detect life extension by metformin (Alfaras et al., 2017; Anisimov et al., 2011) or revealed toxicity that was partially attributed to the metformin overdose (Cabreiro et al., 2013; Martin-Montalvo et al., 2013; Thangthaeng et al., 2017). Strikingly, a dose of 50mM metformin which triggered strongest lifespan extension in a seminal study performed in young *C. elegans*, was moderately toxic when given to middle aged (adulthood day 8) nematodes (Cabreiro et al., 2013; Onken and Driscoll, 2010). We thus came to a conclusion that the benefits and safety of metformin administration to old non-insulin resistant individuals were not sufficiently investigated, contrary to responses of diabetic patients.

Here we used genetic tests, metabolic measurements, stress reporter assays and omics analyses to detect age-specific effects of metformin in *C. elegans* and human primary cells. We found that metformin treatment initiated in late life shortens life span and limits cell survival by aggravating aging associated mitochondrial dysfunction towards respiratory failure. In addition to mitochondrial distortion, old cells failed to enhance the use of glycolysis in response to metformin leading to persistent ATP exhaustion. We found that interventions stabilizing cellular ATP levels, such as ATP repletion and TOR inhibitor rapamycin, alleviate late life metformin toxicity *in vitro* and *in vivo*. We also discovered that early age metformin treatment instigates a range of stress and metabolic adaptations which likely underlay longevity extension by metformin. Importantly, the induction of these favorable responses was strongly impaired in late life. Particularly, we show that early but not late life metformin treatment induces a lipid turnover response similar to dietary restriction (DR). We also found that this early life DR mimetic phenotype is instigated by the triglyceride lipolysis pathway regulated by the protein kinase A and not by AMP activated protein kinase (AMPK) pathway, as suggested previously. Subsequently, we showed that metformin treatment restricted to early adulthood is sufficient for life extension, strengthening the key role of early life stress and metabolic adaptations in longevity benefits of metformin. Finally, we demonstrate that *daf-2(e1370)* mutants, carrying diabetes-like insufficiency of the *C. elegans* insulin receptor, are resilient to late life metformin toxicity in comparison to age-matched wild type controls, due to improved capacity to sustain ATP synthesis during old age metformin exposure. Collectively, we uncovered an alarming capability of metformin to induce metabolic failure in non-insulin resistant old subjects which may limit its benefits for non-diabetic elderly humans.

## Results

### Late life metformin treatment is detrimental for longevity

To address the outcomes of metformin treatment at different age, we treated young adult (3 days old, day 1 of adulthood), adult at the age of reproduction decline (day 4 of adulthood), middle aged (day 8 of adulthood) and old (day 10 of adulthood) wild type *C. elegans* worms with different doses of metformin – 10mM, 25mM and 50mM. 50mM metformin is the common dose used to induce lifespan extension in *C. elegans* while 10mM is the lowest dose linked to reproducible life extension in this model in previous reports (Cabreiro et al., 2013; Onken and Driscoll, 2010; Pryor et al., 2019). We found that metformin treatment started at young age (days 1 and 4 of adulthood) extended lifespan of nematodes at all doses used (Figure 1A and Figure S1A). Within treatment initiated on day 8 of adulthood, the doses of 50mM and 25mM metformin reduced median lifespan but extended maximal lifespan consistent with previous observations (Cabreiro et al., 2013) while 10mM dose was longevity-extending with no detrimental effects (Figure S1B). Strikingly, on day 10 of adulthood metformin was toxic at all doses used with a large proportion of drug-exposed animals dying within first 24 hours of treatment (Figure 1B). Our first experiments in nematodes thus revealed an evident age-dependent decrease in metformin tolerance which culminated in late life toxicity of all metformin doses tested, indicating possible safety risks of late life metformin administration.

**Figure 1.**
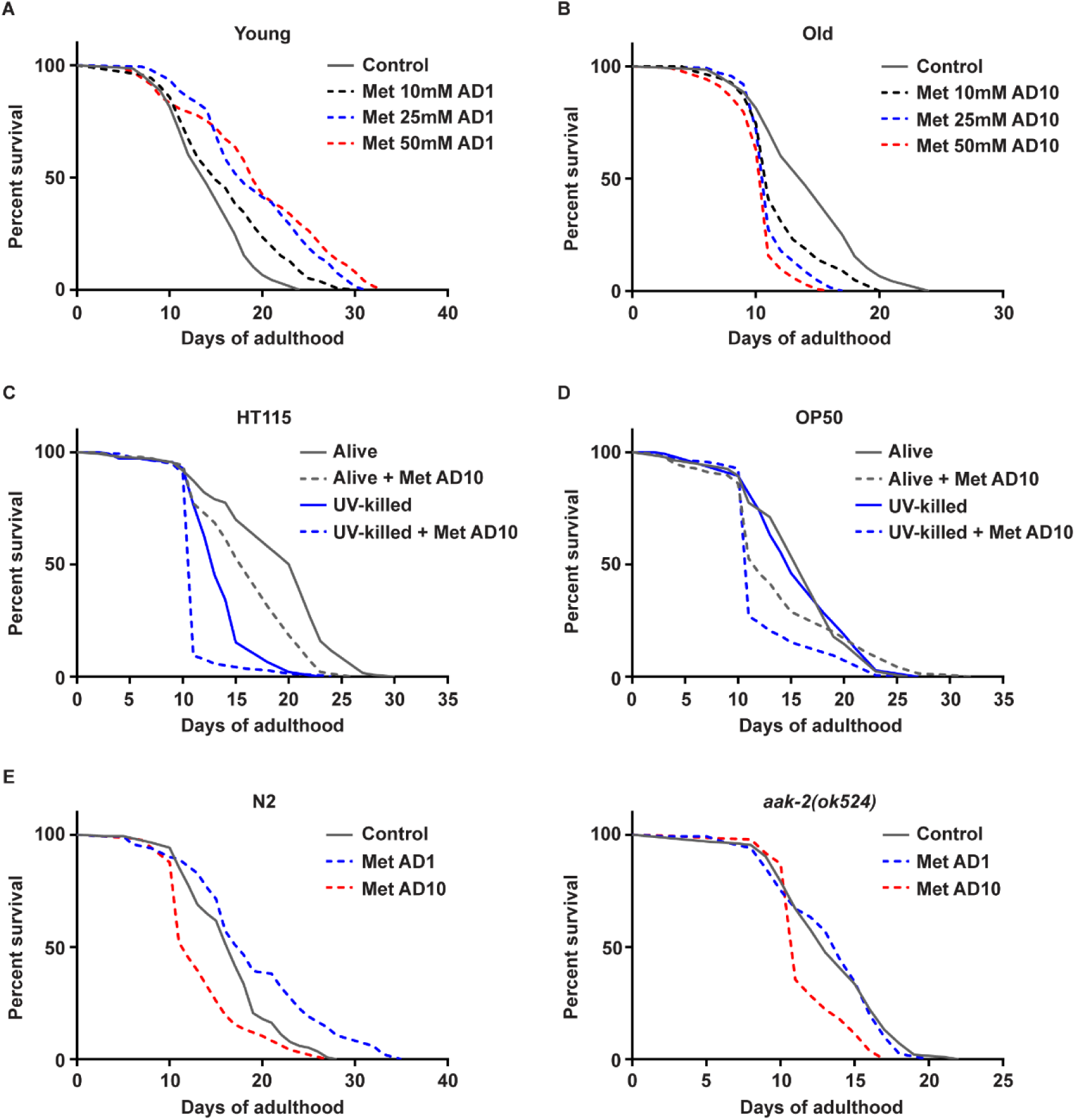
Metformin treatment initiated in late life exerts toxicity and limits survival independently of AMPK and microbial changes. Wild type (WT, N2 Bristol strain) nematodes were treated with indicated doses of metformin (Met) on day 1 (**A**) and day 10 (**B**) of adulthood (AD1 and AD10 respectively), survival was scored daily. WT nematodes were grown on alive and UV-killed HT115 (**C**) and OP50 (**D**) *E. coli* and treated with 50mM metformin on AD10. Survival was scored daily. WT (**E, left**) and AMPK deficient (**E, right**) nematodes were treated with 50mM metformin on AD1 and AD10, survival was scored daily. Significance was measured by log-rank test, n≥100 in all cases, the exact n numbers and statistical values for all panels are presented in Table S1. Each experiment was repeated ≥3 times; one representative result is shown in all cases.

### Late life metformin toxicity is independent of microbiome

To understand the mechanism of late life metformin toxicity, we first addressed known pathways regulating lifespan extension by this compound. The pro-longevity effect of metformin in young *C. elegans* nematodes was previously linked to changes of microbial metabolism induced by this drug (Cabreiro et al., 2013). To test if old age metformin toxicity relied on similar microbiome alterations we treated worms with metformin in the presence of living and UV-killed OP50 (metformin sensitive) and HT115 (metformin-resistant) *E. coli* strains. Old age metformin toxicity developed regardless of the bacterial viability and/or strain (Figure 1C and D) suggesting that late life metformin intolerance is independent of previously uncovered microbiome changes. Interestingly, the baseline survival of nematodes differed between UV killed OP50 and HT115 diets in line with recently reported dependence of nematode physiological behaviors on the bacterial source (Revtovich et al., 2019).

### AMPK is not required for late life toxicity of metformin

Another component essential for young age life-extending effect of metformin is AMP-activated protein kinase (AMPK); particularly metformin failed to promote longevity in nematodes lacking AMPK orthologue AAK-2 (Onken and Driscoll, 2010). In order to probe the requirement of AMPK for old age toxicity of metformin we treated young and old wild type and *aak-2(ok524)* mutant animals with this drug. Consistent with previous reports, early life metformin treatment was unable to induce lifespan extension in *aak-2* deficient worms (Figure 1E); at the same time late life metformin toxicity did develop in mutant animals, indicating that life shortening induced by metformin at old age is not executed by AMPK.

### Metformin toxicity is triggered by mitochondrial impairments

One of metformin’s primary functions is to inhibit complex I of the mitochondrial electron transport chain (ETC), affecting mitochondrial membrane potential and ATP production (Andrzejewski et al., 2014; Cameron et al., 2018; Wheaton et al., 2014). Growth inhibition by metformin was previously linked to impaired mitochondrial respiration in nematodes and mammalian cells (Wu et al., 2016). Additionally, the accumulation of damaged and dysfunctional mitochondria, which may enhance the negative impact of ETC complex I inhibitors on cell survival, is one of the best characterized hallmarks of aging (Bratic and Larsson, 2013; Bratic and Trifunovic, 2010; Cellerino and Ori, 2017; Sun et al., 2016; Taylor and Dillin, 2011).

Mitochondrial deterioration comparable to aging occurs prematurely in mutant animals, defective in mitochondrial biogenesis and quality control (Sun et al., 2016; Trifunovic et al., 2004). To test if metformin toxicity during aging (and metformin toxicity in general) is driven by accumulation of mitochondrial impairments, we obtained mutants harboring deficiencies of diverse mitochondrial homeostasis pathways: mitochondrial unfolded protein response (*atfs-1(gk3094)*) (Nargund et al., 2012), mitochondrial biogenesis (*skn-1(zj15)*) (Palikaras et al., 2015), mitochondrial respiration (*isp-1(qm150)*) (Feng et al., 2001) and mitochondrial protein quality control (*ubl-5(ok3389)*) (Benedetti et al., 2006), and treated these animals with metformin along with wild type counterparts. A combination of metformin with congenital mitochondrial impairments led to an early life onset of metformin toxicity in all mutant backgrounds tested (including normally long-lived *isp-1(qm150)* mutants) clearly linking metformin intolerance to elevated abundance of dysfunctional mitochondria (Figure 2A-C, Figure S2A). The same effect (premature onset of metformin toxicity at young age) was observed in nematodes and human primary cells incubated with mitochondrial uncoupling agent carbonyl cyanide-*p*-trifluoromethoxyphenylhydrazone (FCCP) along with metformin administration (Figure 2D-E and Figure S2B). Of note, congenital mitochondrial respiration defects were recently found to limit the life extending interplay between metformin and the microbiome (Pryor et al., 2019) in line with the key role of mitochondrial integrity in diverse longevity benefits of metformin. To test if mitophagy and/or mitochondrial unfolded protein response (UPR MT) - the protective pathways responding to mitochondrial failure and known to deteriorate during aging (Sun et al., 2016), were directly induced by metformin we measured the abundance of mitochondrial proteins and the expression of *hsp-6::gfp* transgene (UPR MT reporter) in young and old animals treated with this drug. At both ages metformin administration didn’t lead to either elevated expression of GFP or reduction of mitochondrial protein levels (Figure S3A-C), indicating that mitophagy and UPR MT are not prominently triggered by metformin and likely play no role in immediate cellular adaptation to metformin-induced effects. Of note, lack of UPR MT induction by metformin has previously been reported by an independent group (De Haes et al., 2014). We thus show that the early onset of metformin toxicity in mitochondrial mutants is likely driven by the accumulation of mitochondrial damages prior to metformin administration, comparable to what occurs during aging.

**Figure 2.**
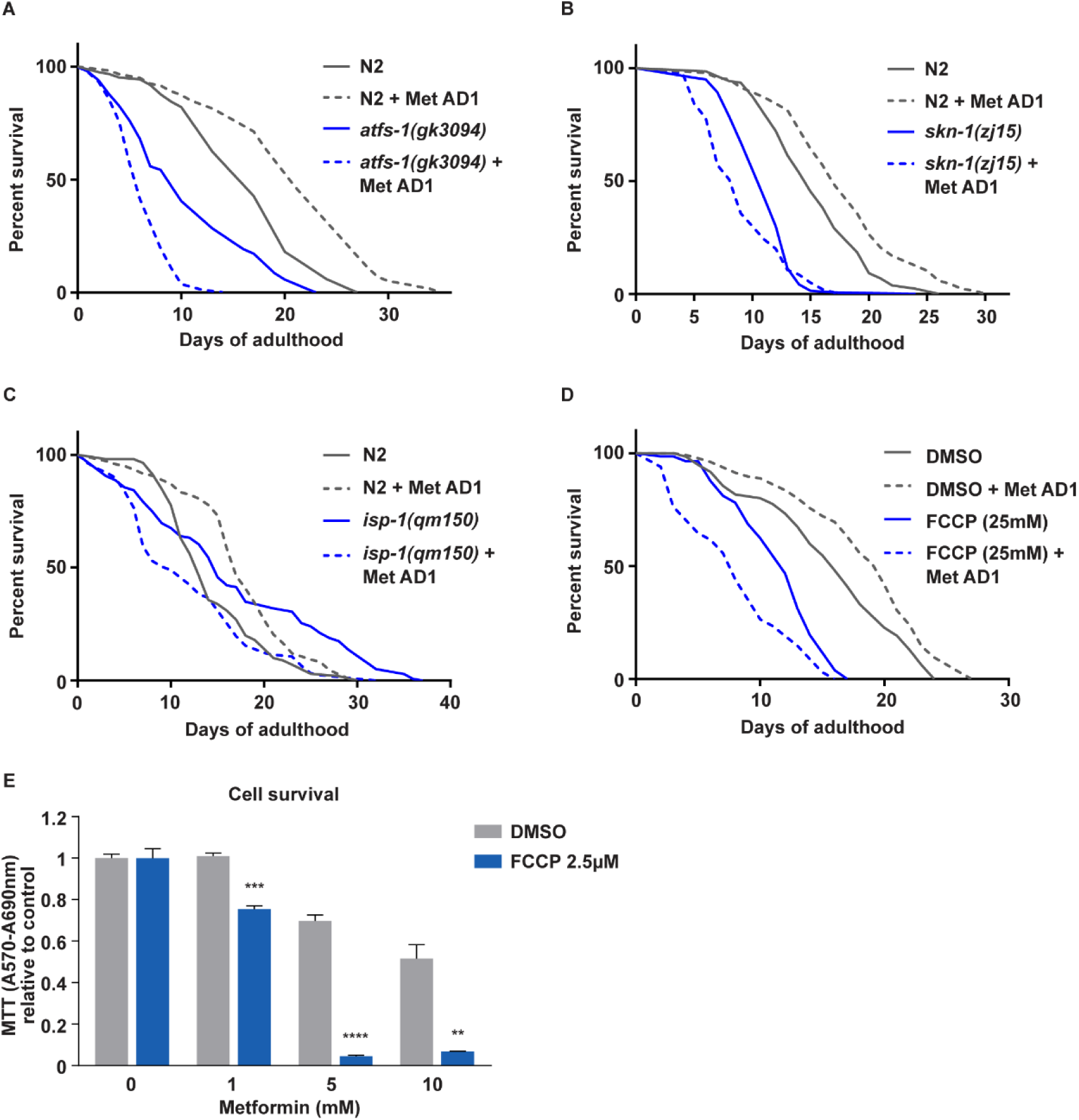
Mitochondrial impairments pre-dispose nematodes and human cells to metformin toxicity. *atfs-1(gk3094)* (**A**), *skn-1(zj15)* (**B**) and *isp-1(qm150)* (**C**) nematodes were exposed to 50mM metformin on adulthood day 1 (AD1) along with age-matched WT animals, survival was scored daily. AD1 WT nematodes (**D**) and early passage (population doubling, PD37) human primary fibroblasts (**E**) were co-treated with FCCP and metformin (50mM in worms), survival was measured daily in worms and after 24h of metformin treatment in cells; DMSO was used as a vehicle control for FCCP in both cells and worms; in **E** all values are relative to a respective untreated (no metformin) condition (separately for vehicle and FCCP). For **A-D**, significance was measured by log-rank test; all n numbers (n≥100 in all cases) and statistical values are presented in Table S1. For **E** n=3, mean and SEM are presented, two-tailed unpaired t-test was used for the statistical analysis, all statistical values are shown in Table S2; ** p<0,01; *** p<0,001; **** p<0,0001.

### Metformin toxicity associates with ATP exhaustion and is alleviated by ATP repletion

To measure direct age-specific effects of metformin on mitochondrial performance and to test the conservation of our findings in humans, we analyzed the impact of metformin treatment on homeostasis of early passage (young) and late passage (old, replicative senescent) primary human skin fibroblasts. The replicative senescence model was chosen because of its high relevance for normal human aging as senescent cells accumulate in aging tissues. *In vitro* aging was performed according to standard procedures demonstrated to yield cells carrying key hallmarks of aging (Tigges et al., 2014), and aging-associated mitochondrial decline of late passage cells was verified by the Seahorse analysis (Figure 3C-D, Figure S4A and C). We found that, at high doses, metformin was toxic to both young and old cells but old cells (similar to old nematodes) showed a much stronger viability decline and succumbed to toxicity already at lower doses of metformin (Figure 3A-B). The oxygen consumption rate (OCR) measurements demonstrated a significant effect of metformin on basal respiration in young and old cells (Figure 3C, Figure S4A), while inhibition of mitochondrial ATP synthesis by metformin was stronger in old fibroblasts (Figure S4C). In addition, only old cells showed a decrease of maximal respiration in response to metformin (Figure 3D, Figure S4A) consistent with a stronger negative impact of this drug on mitochondria of old cells. Interestingly, the extracellular acidification rate (ECAR) was potently increased in young metformin treated fibroblasts (Figure 3E, Figure S4B) in line with their elevated reliance on glycolysis in response to mitochondrial insufficiency triggered by metformin. This adaptive increase of glycolysis was markedly reduced in old metformin exposed cells (Figure 3E, Figure S4B), depriving these cells, in combination with the stronger mitochondrial hindrance by metformin, of effective pathways of ATP synthesis. Subsequent determination of cellular ATP content indeed showed a stronger decline of ATP levels in old metformin treated cells (Figure 3F). We also detected a stronger distortion of the mitochondrial membrane potential in old compared to young metformin exposed fibroblasts (Figure 3G), in line with a more potent mitochondrial decline observed in old cells via oxygen consumption analysis.

**Figure 3.**
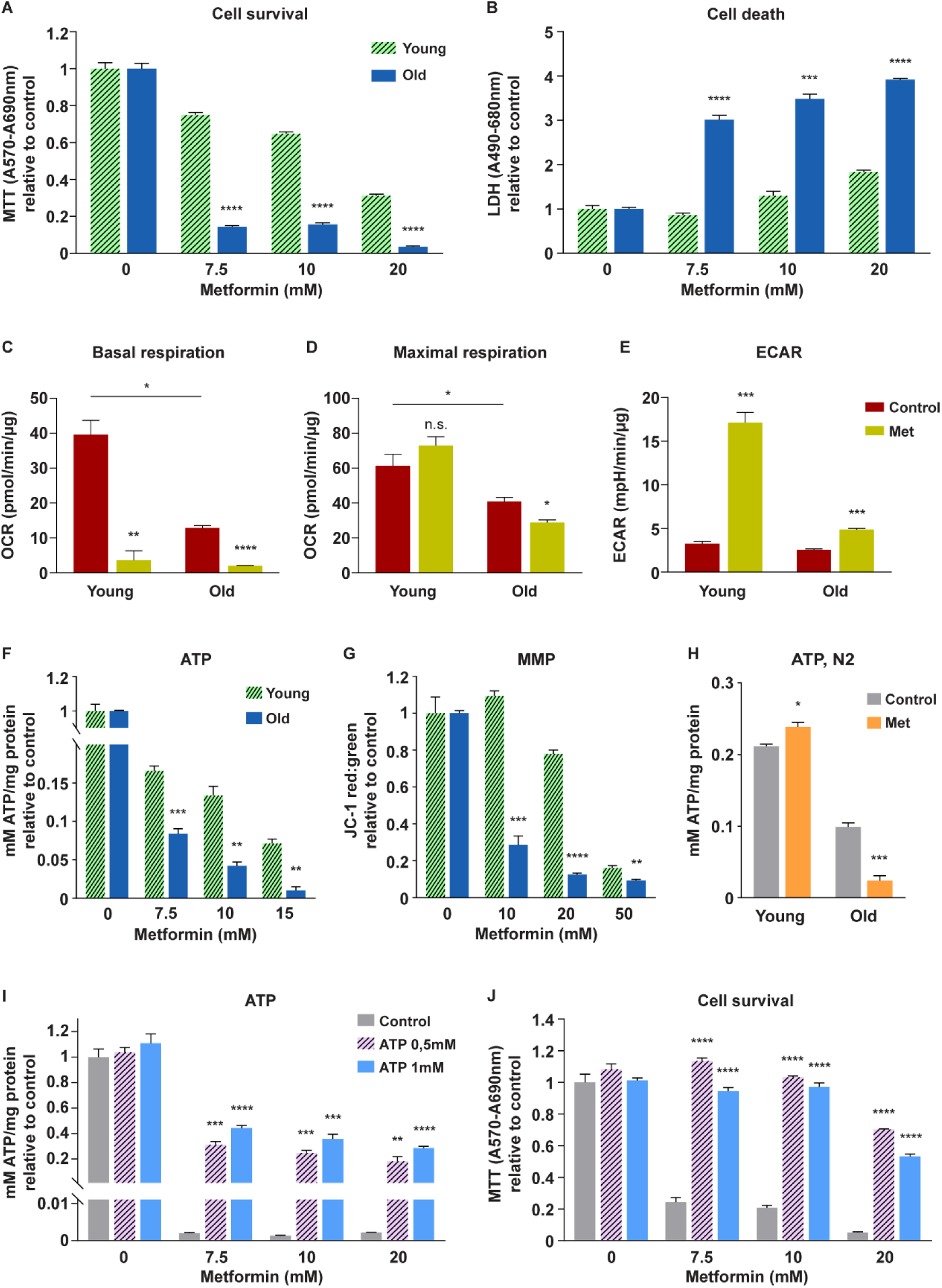
Metformin toxicity associates with loss of mitochondrial homeostasis and ATP exhaustion in human cells and nematodes. Young (PD36) and old (PD61) human skin fibroblasts were treated with metformin for 15 (**F, G**) or 20 (**A, B**) hours; cell survival (**A,** MTT assay), cell death (**B,** LDH assay), mitochondrial membrane potential (**G,** JC-1 assay) and ATP content (**F**) were measured; all values are relative to the untreated control of a given age. (**C**–**E**) Young (PD29) and old (PD61) primary human skin fibroblasts were treated with metformin for 16h, then washed and pre-incubated with media containing high glucose (10mM), glutamine (2mM) and pyruvate (1mM); continuous oxygen consumption rates (OCR) and extracellular acidification rates (ECAR) were recorded following injection of oligomycin (1μM), FCCP (3μM), antimycin-A (0,5μM) and rotenone (0,5μM) (shown in Fig S4). The recorded values were normalized to protein content, basal respiration (**C**), maximal respiration (**D**) and basal ECAR (**E**) were quantified. (**H**) Young (adulthood day 1) and old (adulthood day 10) wild type nematodes were treated with 50mM metformin, whole organism ATP levels were measured after 36h. Pre-senescent (PD44) fibroblasts were treated with metformin for 24h in presence or absence of ATP; ATP content (**I**) and cell survival (**J**) were measured; all values are relative to the untreated control (no metformin, no ATP) for each assay. For **H** n=100, for all other panels n=3, mean and SEM are presented, two-tailed unpaired t-test was used for the statistical analysis, all statistical values are presented in Table S2; * p<0,05; ** p<0,01; *** p<0,001; **** p<0,0001.

We next measured the effect of metformin on organismal ATP content in nematodes. Metformin treatment of young animals didn’t cause negative changes of systemic ATP levels (Figure 3H), consistent with reduced mitochondrial impairments and intact metabolic adaptability at young age. Strikingly, old animals showed a strong reduction of baseline ATP levels, compared to untreated young animals, followed by a further 76% decline of ATP content induced by metformin (Figure 3H, Table S2). These results are consistent with known mitochondrial deterioration and reduced energetics of old animals (and cells) (Brys et al., 2010; Drew et al., 2003), and suggest, along with above reported age-specific metabolic phenotypes, that metformin toxicity may be linked to a failure of old cells to maintain ATP synthesis during metformin treatment, leading to a decline of ATP content down to levels incompatible with cell viability. To test this hypothesis we asked if ectopic ATP supplementation would rescue metformin toxicity. Providing ATP to animals is not feasible due to insufficient bioavailability upon oral supplementation (Arts et al., 2012), we were however able to supplement ATP to fibroblasts in culture using millimolar concentrations as previously described (1978). Ectopic ATP repletion alleviated cell death, ATP exhaustion and loss of mitochondrial membrane potential induced by metformin (Figure 3I-J, Figure S5A-D) and by another complex I inhibitor rotenone (Figure S6A-C) linking viability loss upon treatment with ETC complex I inhibitors to alterations in energy homeostasis. The rescue of the mitochondrial membrane potential by ATP repletion was consistent with the previously reported ability of the mitochondrial ATP synthase to restore membrane potential under conditions of respiratory failure by consuming ATP (Chinopoulos and Adam-Vizi, 2010).

### Metformin toxicity is suspended in nematodes carrying insulin receptor insufficiency

Because metformin intolerance, comparable to our findings, has not been reported in type 2 diabetes patients receiving treatment in late life, we decided to test the old age metformin response of nematodes lacking functional insulin receptor (a molecular mimetic of insulin resistance in type 2 diabetes). For that we chose a well characterized *daf-2(e1370)* strain which carries a temperature sensitive mutation of the *C. elegans* IGF-1/Insulin receptor orthologue DAF-2 (Dorman et al., 1995). At a restrictive temperature of 20°C *daf-2(e1370)* mutants exhibit partial loss of insulin receptor function leading to a variety of phenotypes, partially comparable to diabetic patients (such as enhanced fat deposition) (Kimura et al., 1997; Tissenbaum and Ruvkun, 1998). We next measured survival of *daf-2(e1370)* and wild type nematodes treated with metformin on day 10 of adulthood (AD10) and observed a striking resilience of the mutants to metformin killing at this age (Figure 4A). Systemic ATP measurements revealed a moderate reduction of baseline ATP synthesis in young *daf-2(e1370)* mutants (Figure 4C) consistent with previous reports (Brys et al., 2010; Palikaras et al., 2015). Interestingly neither aging nor metformin had a negative impact on ATP levels in AD10 *daf-2(e1370)* animals, contrary to the response of wild type controls (Figure 4C), associating the resilience of the mutants to metformin toxicity with their ability to sustain ATP synthesis during late life metformin treatment. Of note, the enhanced competence of *daf-2(e1370)* mutants in ATP synthesis during late life is also consistent with previous reports (Brys et al., 2010; Palikaras et al., 2015). Importantly, the resistance of *daf-2(e1370)* animals to metformin wasn’t infinite and mutant nematodes exposed to the drug on adulthood day 21 (AD21) showed levels of metformin toxicity comparable to AD10 wild type controls (Figure S7A), indicating that the insulin receptor itself is not required for the induction of late life metformin toxicity. In summary, these data reiterate the key role of ATP exhaustion in late life metformin toxicity and indicate that inhibition of insulin receptor signaling, comparable to insulin resistance in diabetes, provides extended protection from late life metformin intolerance.

**Figure 4.**
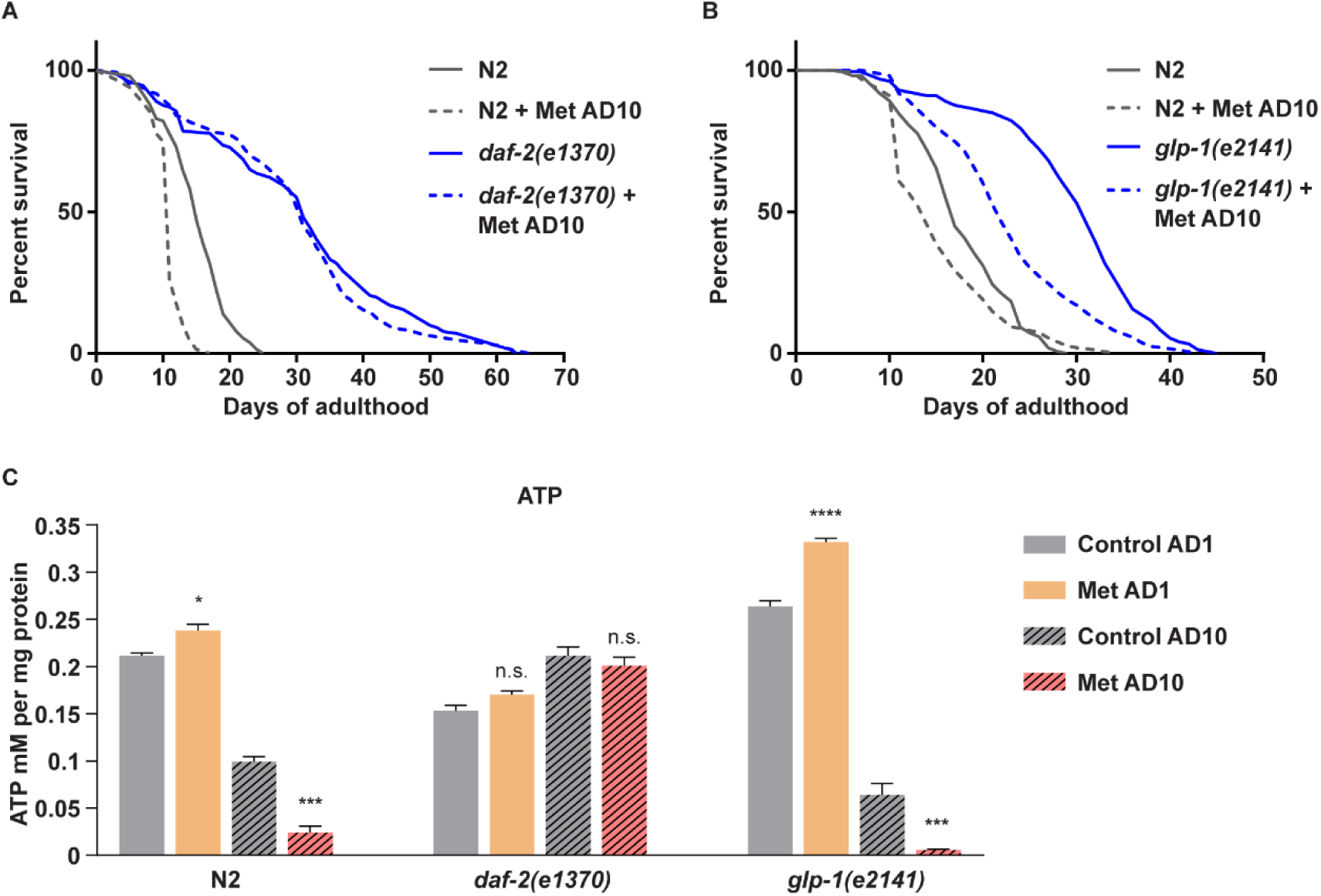
Insulin receptor deficiency suspends late life metformin toxicity in *C. elegans* by preventing metformin-triggered ATP exhaustion. *daf-2(e1370)* (**A**) and *glp-1(e2141)* (**B**) temperature sensitive mutants and corresponding wild type (N2 Bristol strain) control animals were grown at 20°C and 25°C respectively to achieve inactivation of *daf-2* and *glp-1* pathways. All animals were treated with 50mM metformin (Met) on adulthood day 10 (AD10), survival was scored daily. (**C**) Whole organism ATP levels were measured in wild type, *daf-2(e1370)* and *glp-1(e2141)* mutant animals treated with 50mM metformin on AD1 and AD10, the measurement was performed after 36h of metformin exposure. For **A-B** significance was measured by log-rank test; n numbers (n≥100 in all cases) and statistical values are presented in Table S1. For **C** n=100 for each condition, mean and SEM are presented, two-tailed unpaired t-test was used for the statistical analysis, all statistical values are presented in Table S2; * p<0,05; *** p<0,001; **** p<0,0001.

### Metformin resilience of insulin receptor mutants is not due to their slower rate of aging

The *daf-2(e1370)* mutants are known to be long-lived in comparison to wild type animals (Dorman et al., 1995; Kimura et al., 1997; Tissenbaum and Ruvkun, 1998). To ensure that the prolonged metformin resilience of *daf-2(e1370)* animals is not a simple consequence of their slower rate of aging, we selected a different comparably long-lived strain - germline-deprived *glp-1(e2141)* mutants (Lapierre et al., 2011), and treated these animals with metformin on day 10 of adulthood (AD10). Unlike *daf-2(e1370)* nematodes, *glp-1(e2141)* worms succumbed to AD10 metformin killing similar to wild type controls (Figure 4B). They also showed both aging-triggered and metformin-aggravated declines of ATP levels, unlike AD10 *daf-2(e1370)* mutants and similar to wild type controls (Figure 4C). We thus determined that the ability to sustain ATP synthesis in late life and the corresponding metformin resilience are not common phenotypes of all long-lived mutants but rather a specific feature of insulin receptor deficient animals.

### Metformin resilience of insulin receptor mutants depends on sustained ATP synthesis in late life regulated by DAF-16/FOXO

The exact mechanism behind the enhanced metabolic competence of *daf-2(e1370)* mutants in late life remains to be elucidated. However, similar to most of *daf-2(e1370)* beneficial phenotypes, this capacity may depend on the FOXO transcription factor DAF-16 (Kenyon, 2011; Palikaras et al., 2015). Consistently, *daf-2(e1370); daf-16(mu86)* double mutants didn’t show metformin resilience on adulthood day 10, unlike *daf-2(e1370)* single mutants and similar to wild type controls (Figure S7B). The double mutants also showed a clear decline of ATP synthesis upon AD10 metformin treatment (Figure S7C) in line with the requirement of DAF-16 for metabolic resilience of the *daf-2(e1370)* genetic background. Because DAF-16 is an important mediator of multiple stress adaptations (Kenyon, 2011), we tested if the response to metformin required DAF-16 directly by measuring nuclear translocation of the DAF-16::GFP fusion protein during metformin or heat shock treatment (used as positive control). While DAF-16 was clearly translocating into the nucleus in response to heat stress, little translocation (above negative control values) could be observed in the course of metformin treatment regardless of age (Figure S8A-C), suggesting that metformin resilience of *daf-2(e1370)* mutants is not driven by the direct involvement of DAF-16 in stress adaptations during metformin treatment.

The capability of *daf-2(e1370)* animals to sustain stable ATP synthesis at old age has previously been linked to elevated mitochondrial turnover induced by life-long subtitle metabolic stress, leading to improved mitochondrial quality in late life (Brys et al., 2010; Palikaras et al., 2015). Our proteomics measurements indeed revealed reduced baseline levels of mitochondrial proteins in *daf-2(e1370)* mutants compared to wild type animals (Figure S7D) consistent with elevated mitochondrial turnover. Interestingly, mitochondrial protein levels of *daf-2(e1370); daf-16(mu86)* double mutants were similar to values observed in wild type animals (Figure S7D), suggesting that mitochondrial replenishing triggered by insulin receptor deficiency is inhibited in the absence of DAF-16. Consistent with the key role of DAF-16 in supporting metabolic homeostasis of *daf-2(e1370)* mutants, the *daf-2(e1370); daf-16(mu86)* double mutants demonstrated strong impairments of baseline ATP synthesis already at young age (Figure S7C).

### Metformin rapidly activates diverse longevity assurance pathways in young but not old animals

To test if other metformin responses, in addition to ATP synthesis, become altered in late life we performed an unbiased proteomics analysis of animals treated with metformin on days 1 and 10 of adulthood for 24 and 48 hours, and compared these to age- and time point-matched untreated controls. The abundance of only a restricted proportion of protein groups was affected by metformin at both ages and at both times of treatment (seen by comparing numbers of altered proteins (circled) and non-altered proteins (outer box) in Figures 5A and S9A), highlighting the specificity of the response elicited by the drug treatment. By comparing the protein expression profiles obtained (4572 protein groups were quantified in total, Table S3), we determined that global responses to metformin at young and old age were largely distinct, as indicated by a limited overlap between commonly affected proteins at both 24 and 48 hours of treatment (Figure 5A and S9A), and a modest correlation between induced protein changes (Figure 5B and S9B). Interestingly, the similarity between young and old responses was stronger at 48 hours of treatment, which could be explained either by a delay of old animals in responding to the drug or by an enrichment of the 48h old age metformin-exposed sample with animals that were most competent in their adaptation to metformin treatment (based on our survival analyses a significant proportion of metformin sensitive old animals died between 24 and 48h of treatment).

**Figure 5.**
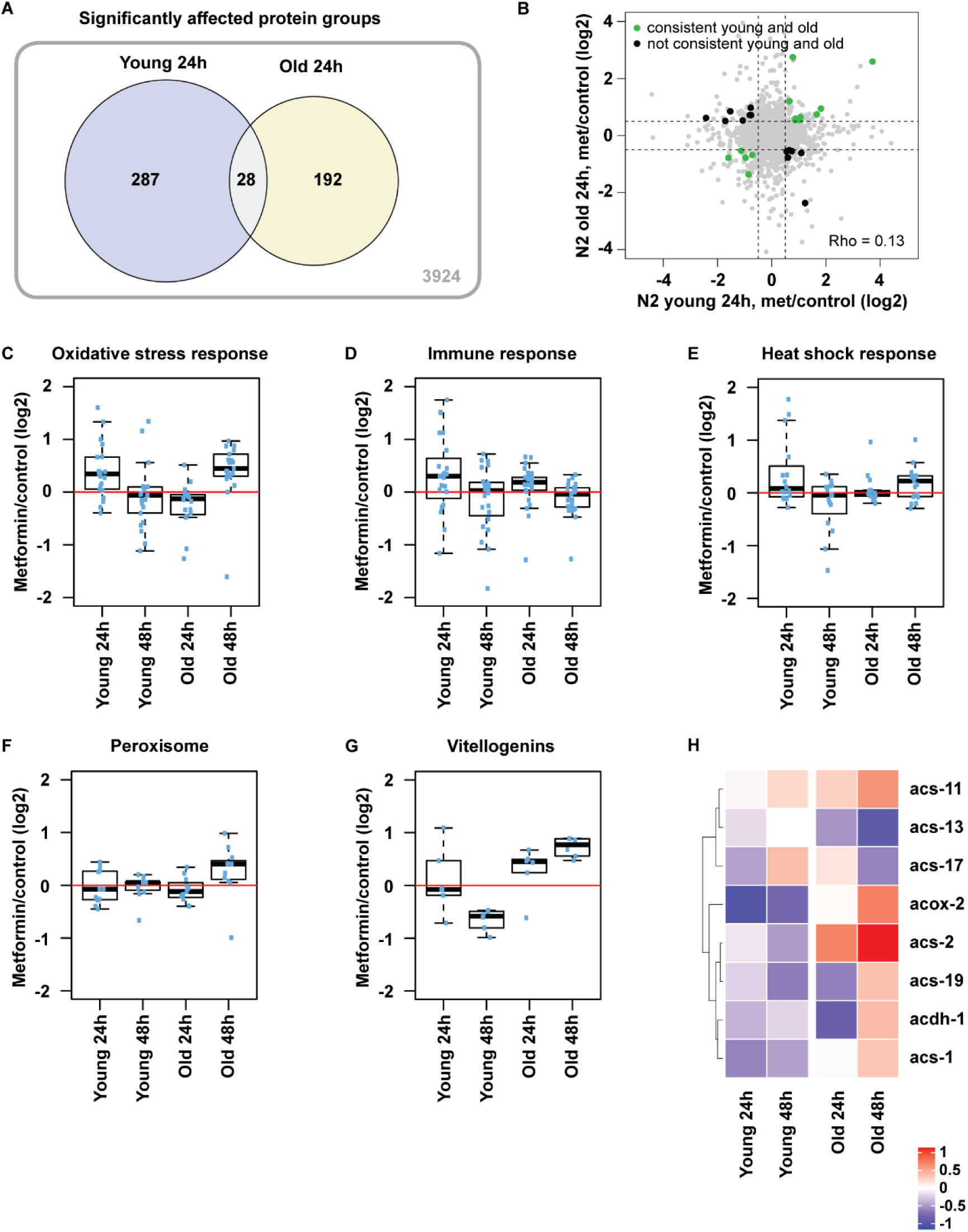
The induction of longevity assurance pathways and metabolic adaptations by metformin is age-dependent. Wild type worms were treated with 50mM metformin on adulthood day 1 (young) and 10 (old). (**A**) Venn diagram showing overlap between significantly altered proteins after 24h of early and late life metformin treatment is shown; the circled values include all proteins with absolute log2 fold change above 0.5 and Q value below 0.25; the number in the outside box indicates non-regulated proteins detected in both young and old samples collected at 24h; fold changes were calculated against age- and time point matched untreated controls. Three independent pools of worms were analyzed for each sample group. (**B**) Scatter plot comparing log2 fold changes of individual proteins (shown as dots) after 24h of metformin treatment at young and old age is shown. Proteins significantly regulated at both ages are highlighted with color: the green-highlighted proteins are consistently regulated at both ages while black-highlighted ones are regulated in the opposite direction between young and old age. The correlation coefficient (Rho) between young and old responses taking into account all regulated proteins is shown. Boxplots showing fold changes for selected proteins involved in oxidative stress response (**C**), immune response (**D**), heat stress response (**E**) as well as peroxisomal proteins (**F**) and vitellogenins (**G**) are presented. The median fold change of the proteins belonging to each group is shown as a bold line, the upper and lower limits of the boxplot indicate the first and third quartile, respectively, and whiskers extend 1.5 times the interquartile range from the limits of the box. (**H**) Heatmaps of selected proteins implicated in lipid metabolism are shown, only fold changes with significance in at least one age/treatment combination are depicted. n≥500 and 3 replicas were measured for each condition. The list and complete data of proteins used for each boxplot and the heatmap are reported in Table S3.

By analyzing pathways regulated by metformin at distinct age points, we detected a rapid activation of adaptive stress responses and longevity assurance pathways such as downregulation of the ribosome (MacInnes, 2016) and induction of oxidative stress response (Honda and Honda, 1999), immune response (Xia et al., 2019) and heat stress response (Baldi et al., 2017) in young animals (Figure 5C-E and S9C). General autophagy (Hansen et al., 2018) was also comparably regulated albeit to a smaller extent (Figure S9D). These data are consistent with previous studies suggesting that metformin extends life span by inducing stress adaptations such as oxidative stress response (De Haes et al., 2014; Onken and Driscoll, 2010). Unlike previous reports, however, our data demonstrates for the first time that metformin simultaneously triggers a complex array of diverse stress responses which likely all contribute to life extension by this drug. Strikingly, the induction of stress adaptations by metformin was either inhibited or delayed in old animals (Figure 5C-E, Figure S9C-D) suggesting that, in addition to metformin-triggered ATP exhaustion, also the longevity assurance component of the metformin response is impaired in late life.

To investigate the age specificity of metformin-triggered stress responses by an independent method, we measured the induction of general autophagy by metformin in a transgenic strain expressing the LGG-1::mCherry fusion protein (Gosai et al., 2010). LGG-1 is a *C. elegans* LC3 orthologue that gets recruited to autophagosome membranes during the autophagy process, giving rise (in a reporter setting) to distinct puncta which can be quantified microscopically (Figure S10A). Autophagy was chosen for the validation test because of its known antagonistic effect on longevity depending on age (Wilhelm et al., 2017) and mitochondrial integrity (Zhou et al., 2019). By treating young and old transgenic animals with metformin, we could indeed observe a substantial induction of autophagosome formation in young but not old metformin-exposed animals (Figure S10B-C) with effects size comparable to the proteomics data (Figure S9D). Interestingly, the absolute numbers of autophagy puncta in control untreated animals were significantly higher at old age (Figure S10D-E) consistent with the previously observed inhibition of autophagy flux during late life (Wilhelm et al., 2017). The specificity of observed puncta to the autophagy process was confirmed by exposing transgenic animals to RNAi against key autophagy mediator beclin 1 (*bec-1*) (Figure S10F).

In summary, we uncovered that metformin treatment exerts an array of stress adaptation responses in young animals while old animals fail to activate these signals to a comparable extent in line with previous hints of impaired stress resilience at old age (Haigis and Yankner, 2010). To probe the contribution of early life adaptive events to the pro-longevity effect of metformin we exposed nematodes to the drug during early adulthood and indeed found that metformin treatment restricted to young age is sufficient to confer *in vivo* life span benefits (Figure S10G).

### Old age specific metformin response is enriched in distinct mediators of lipid metabolism

We next screened the proteomics data for pathways activated by metformin predominantly in old animals. We found that peroxisomal components along with mitochondrial and peroxisomal enzymes implicated in fatty acid β-oxidation (such as acs-1, acs-2, acox-2) (Zhang et al., 2011) were upregulated in old stronger than in young animals (Figure 5F, H). Also proteins associated with lipid droplets, serving as key lipid storage units in *C. elegans*, (vitellogenins, dehydrogenases) (Vrablik et al., 2015; Zhang et al., 2012) were upregulated in old while they were downregulated in young metformin treated animals (Figure 5G, Figure S9E). In addition, ribosomal components found to be part of the lipid droplet proteome (Vrablik et al., 2015; Zhang et al., 2012) were downregulated in young but not in old metformin exposed nematodes (Figure S9C). Collectively, our data indicated that metformin treatment leads to an increase in peroxisomal content, which is stronger at old compared to young age, while lipid droplet components are downregulated in young and upregulated in old animals. Interestingly, the downregulation of lipid droplets and the upregulation of peroxisomes by metformin in young animals were recently reported by an independent study (Pryor et al., 2019) which however didn’t address the age specificity of these phenotypes.

### Metformin treatment leads to distinct lipid turnover responses in young and old animals

Because proteomics analysis revealed the age-specific regulation of lipid turnover mediators by metformin, we asked if metformin affects systemic lipid deposition in an age-dependent manner. By performing Oil Red O whole body lipid staining in *C. elegans* we detected a significant decline of lipid levels in young animals exposed to metformin while no such decline could be observed in old animals (Figure 6A, Figure S12A). Importantly, a reduction of whole body lipid content by metformin is consistent with its role as a dietary restriction (DR) mimetic (Onken and Driscoll, 2010) which was recently found to be essential for the longevity extension by this drug (Pryor et al., 2019). Our data in old animals demonstrates that this beneficial DR effect of metformin is abolished during aging.

**Figure 6.**
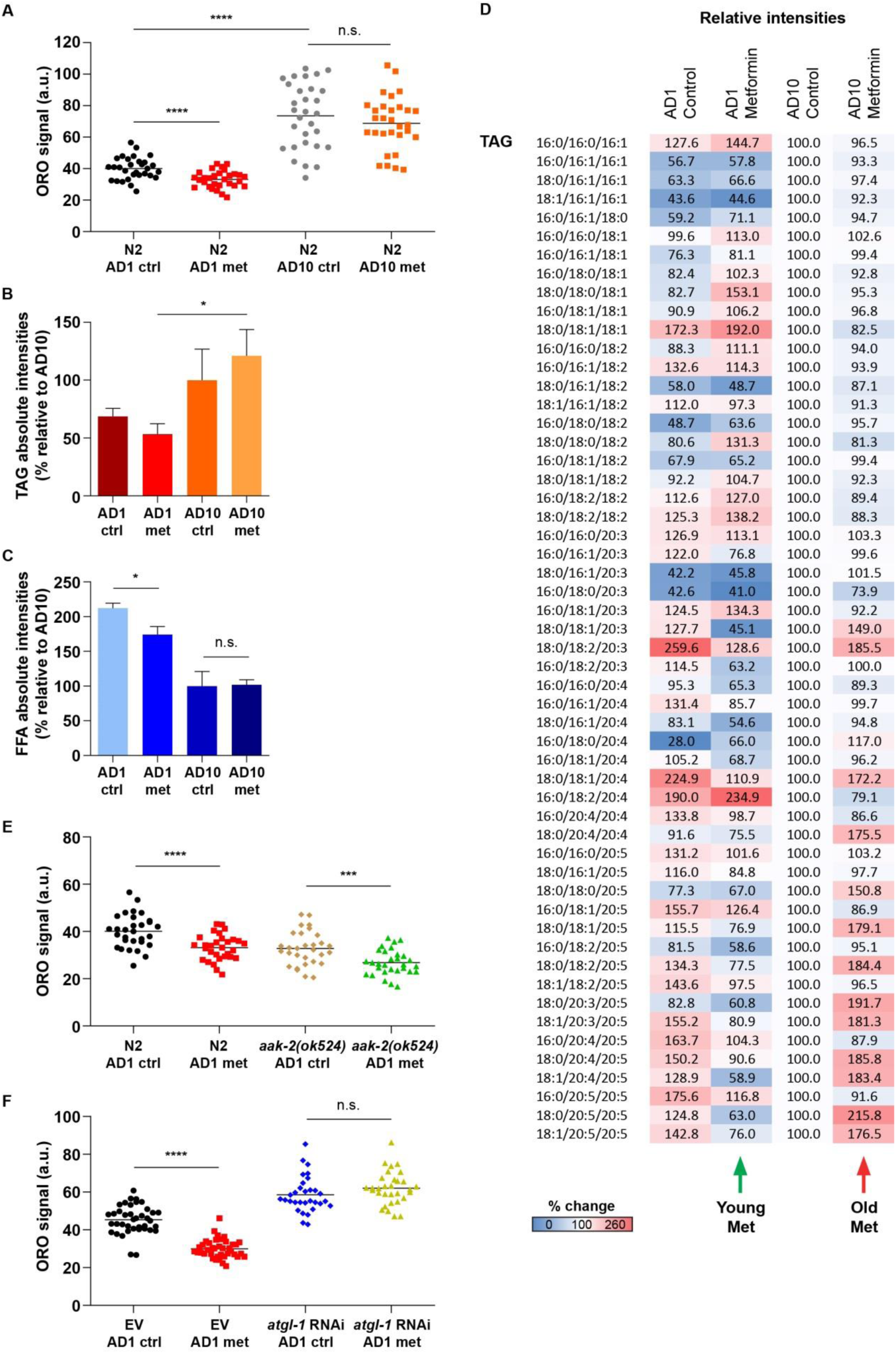
Metformin induces a dietary restriction mimetic lipid turnover response via the protein kinase A pathway in young but not old animals. (**A**) Whole body lipid content was detected by Oil Red O staining in young (adulthood day 1, AD1) and old (adulthood day 10, AD10) worms treated with 50mM metformin for 24h. Wild type (N2 Bristol strain), AMPK deficient (**E**) and *atgl-1* RNAi knock down animals (**F**) were treated with 50mM metformin on adulthood day 1 (AD1) and whole body Oil Red O lipid staining was performed after 24h of treatment; Oil Red O mean grey values (ImageJ) are used as arbitrary units (a.u.) in all cases. Wild type animals were treated with 50mM metformin for 24h on AD1 and AD10; lipids were isolated and analyzed by UPLC-MS/MS, absolute intensities for triglycerides (TAGs) (**B**) and free fatty acids (FAAs) (**C**), and relative intensities for distinct TAGs (**D**) are shown, normalized to the AD10 untreated control in all cases (chosen to highlight the late life specific effects of metformin). Definitions of absolute and relative intensities are provided in the Methods section. For **A** and **E** n=30, for **F** n≥31 and for **B-D** 700 animals were analyzed in 3 replicas for each condition; mean and SEM (for **B**, **C**) are presented, two-tailed unpaired t-test was used for the statistical analysis when applicable, all statistical values are presented in Table S2; * p<0,05; *** p<0,001; **** p<0,0001. All individual values for the lipidomics analysis are summarized in Table S4.

We next performed lipidomics analysis by mass spectrometry in young and old nematodes exposed to metformin to characterize the age-specific lipid turnover phenotype induced by this drug in a greater detail. By comparing lipid abundancies in untreated young and old worms we observed a significant exhaustion of phospholipids (PLs), free fatty acids (FFAs), lysophospholipids and polyunsaturated fatty acid (PUFA)-rich lysophospholipids which occurred due to aging (Figure S13A-B and Table S4). Of note, these observations are consistent with a recent lipidomics report characterizing basal lipid changes in aging *C. elegans* (Gao et al., 2017). Interestingly, the levels of triglycerides (TAGs) were markedly increased in old animals compared to young worms (Figure 6B, D and Table S4), in contrast to other lipid classes measured and reminiscent of the aberrant accumulation of triglycerides observed in human aging (Cree et al., 2004) and upon mitochondrial insufficiency (Vankoningsloo et al., 2006). Importantly, the accumulation of TAGs during aging correlated with the elevated whole body lipid content detected by Oil Red O staining in old untreated nematodes (Figure 6A).

We next compared lipid responses to metformin in young and old animals and found that in young nematodes metformin lowered the abundance of diverse lipid subclasses including free fatty acids and triglycerides, consistent with the DR-like phenotype (Figure 6B-D, Figure S13C-E, Table S4). Strikingly, the levels of triglycerides, including TAGs containing highly unsaturated PUFAs, were further elevated in old metformin exposed animals (Figure 6B, D and Figure S13E), contrary to the response of young nematodes and consistent with the loss of the metformin DR effect at old age. The opposite changes in triglyceride content in young and old animals exposed to metformin were also consistent with the opposite dynamics of lipid droplet-associated proteins observed in these animals by proteomics (Figure 5G, Figure S9C, E): triglycerides are the key components of lipid droplets, and lipid droplet turnover supports the engagement of TAGs during fasting and DR (Lee et al., 2014). Along with the increase in triglycerides and highly unsaturated PUFA-rich triglycerides (Figure S13E), metformin triggered a reduction of long-chain and PUFA-rich lysophospholipids and free fatty acids in old animals (Figure S14A-C and Table S4), similar to aberrant lipid rearrangements observed during persistent metabolic stress (Markel et al., 1985; Nguyen et al., 2017; Steinhauser et al., 2018) and in line with the differential abundance of distinct lipid turnover mediators detected in metformin-exposed old nematodes by proteomics. In summary, old animals failed to develop a DR mimetic lipid turnover phenotype in response to metformin, contrary to young worms, and rather showed an exacerbation of pre-existing aging-associated lipid abnormalities upon metformin treatment. Given the recently reported key importance of lipid turnover, resembling DR, for life extension by metformin (Pryor et al., 2019), the reversal of this DR response during aging likely contributes to the lack of metformin benefits in late life.

### Metformin-triggered lipid changes are driven by distinct mechanisms at young and old age

Previous studies implicated AMPK in lipid changes linked to aging and mitochondrial alterations (Gao et al., 2017; Weir et al., 2017). AMPK has also been linked to a reduction of de-novo lipid synthesis in metformin-exposed hepatocytes, muscle and liver (Boudaba et al., 2018; Collier et al., 2006; Fullerton et al., 2013; Zang et al., 2004). We next asked if lipid transformations triggered by metformin at young or old age were mediated by AMPK. Wild type and AMPK (AAK-2) deficient animals were treated with metformin at young and old age followed by Oil Red O whole body lipid staining. We found that old AMPK deficient animals accumulated additional lipids in response to metformin while no such changes were seen in wild type nematodes (Figure S11A, S12C), in line with an important role of AMPK in mitigating the aberrant lipid accumulation during late life metabolic stress. But curiously, young AMPK deficient nematodes demonstrated an even stronger lipid reduction in response to metformin compared to wild type controls (Figure 6E, S11C and S12B) indicating that AMPK is not the primary instigator of early life metformin DR response but rather plays an inhibitory role in this process.

Because the age-specific lipid turnover induced by metformin affected triglycerides stronger than other lipid species, we next asked if the triglyceride lipolysis pathway regulated by protein kinase A (PKA) in response to fasting (Lee et al., 2014) is the principle driver of early life metformin effect on lipid reserves. By performing Oil Red O whole body lipid staining in control nematodes and in nematodes harboring RNAi knock down of *kin-1* (the *C. elegans* orthologue of PKA) or *atgl-1* (the orthologue of the adipose triglyceride lipase, the key effector of the PKA lipolysis pathway) we found that inactivation of the PKA pathway indeed suppressed the DR-like lipid turnover induced by metformin in early life (Figure 6F, S11B and S12D-E). Interestingly, AMPK was previously shown to directly modulate the activity of ATGL-1 towards a more moderate turnover of triglyceride reserves, and this inhibitory capacity of AMPK was found to be essential for the extended longevity of long-lived *C. elegans* dauer larvae (Narbonne and Roy, 2009). In our tests we observed a stronger decline of lipid content in young AMPK deficient animals exposed to metformin compared to wild type controls (Figure S11C), it is thus feasible that the modulatory effect of AMPK on the PKA lipolysis pathway is behind the known essential role of AMPK in longevity extension by metformin (Onken and Driscoll, 2010), by preventing the untimely lipid exhaustion during early life metformin exposure. Importantly, young animals exposed to metformin in the presence of HT115 *E. coli* (the RNAi vehicle), a condition previously found to be deprived of metformin life extension similar to AMPK mutants (Cabreiro et al., 2013), showed markedly enhanced loss of lipids at 24h of drug treatment compared to OP50 *E. coli* control and similar to AMPK-deficient animals (Figure S11D and S12D-E, empty vector control); this excessive loss of lipids was clearly prevented by the inactivation of ATGL-1, the key triglyceride lipase regulated by AMPK (Kim et al., 2016)(Figure S11D and S12D). Our data thus supports the model of metformin life extension where prevention of untimely and unhealthy lipid loss is as important for improved longevity as the induction of the beneficial DR response, previously linked to metformin induced life prolongation (Onken and Driscoll, 2010; Pryor et al., 2019). We also show that AMPK has age-specific roles in metformin-triggered lipid turnover and that the PKA and not AMPK pathway instigates the DR-like lipid utilization response in young metformin treated animals.

### ATP exhaustion and late life metformin toxicity are alleviated by in vivo rapamycin co-treatment

Because appropriate lipid turnover is essential for supporting the mitochondria with metabolites required for effective ATP synthesis, our molecular analysis of key metformin-modulated pathways further supported the primary role of ATP exhaustion in cellular deterioration triggered by metformin in late life. We next decided to test if accessible *in vivo* interventions which stabilize cellular ATP levels alleviate late life metformin toxicity. One known intervention that stabilizes ATP content under conditions of mitochondrial dysfunction and carbon starvation is TOR inhibitor rapamycin (Thomsson et al., 2005; Zheng et al., 2016). Interestingly TOR inhibition was previously found to be essential to support nematode development and cell survival upon treatment with toxic high doses of metformin (Wu et al., 2016). Previous studies also indicated that rapamycin co-exposure enhances longevity benefits of early life metformin treatment (Strong et al., 2016). We next exposed pre-senescent fibroblasts to metformin in the presence or absence of rapamycin and observed a clear alleviation of metformin toxicity in rapamycin pre-treated cells (Figure 7A and S15A-B) accompanied by blunted ATP exhaustion and reduced loss of mitochondrial membrane potential in these cells (Figure 7B-C and S15C-D). To test if rapamycin co-exposure protects against late life metformin toxicity also *in vivo*, we pre-treated nematodes with rapamycin from adulthood day 8 (AD8) followed by metformin treatment on adulthood day 10. Consistent with our cell culture observations, rapamycin co-treatment significantly alleviated late life metformin intolerance as seen by improvement of both median and maximal life spans in rapamycin/metformin co-exposed animals (Figure 7D). These *in vivo* findings confirm the decisive role of ATP exhaustion in late life metformin toxicity and suggest that interventions capable of stabilizing cellular ATP content, such as rapamycin administration, could be utilized for prevention of metformin intolerance during aging. The incomplete reversal of late life metformin toxicity by rapamycin is in line with the limited capacity of this drug to preserve cellular ATP levels.

**Figure 7.**
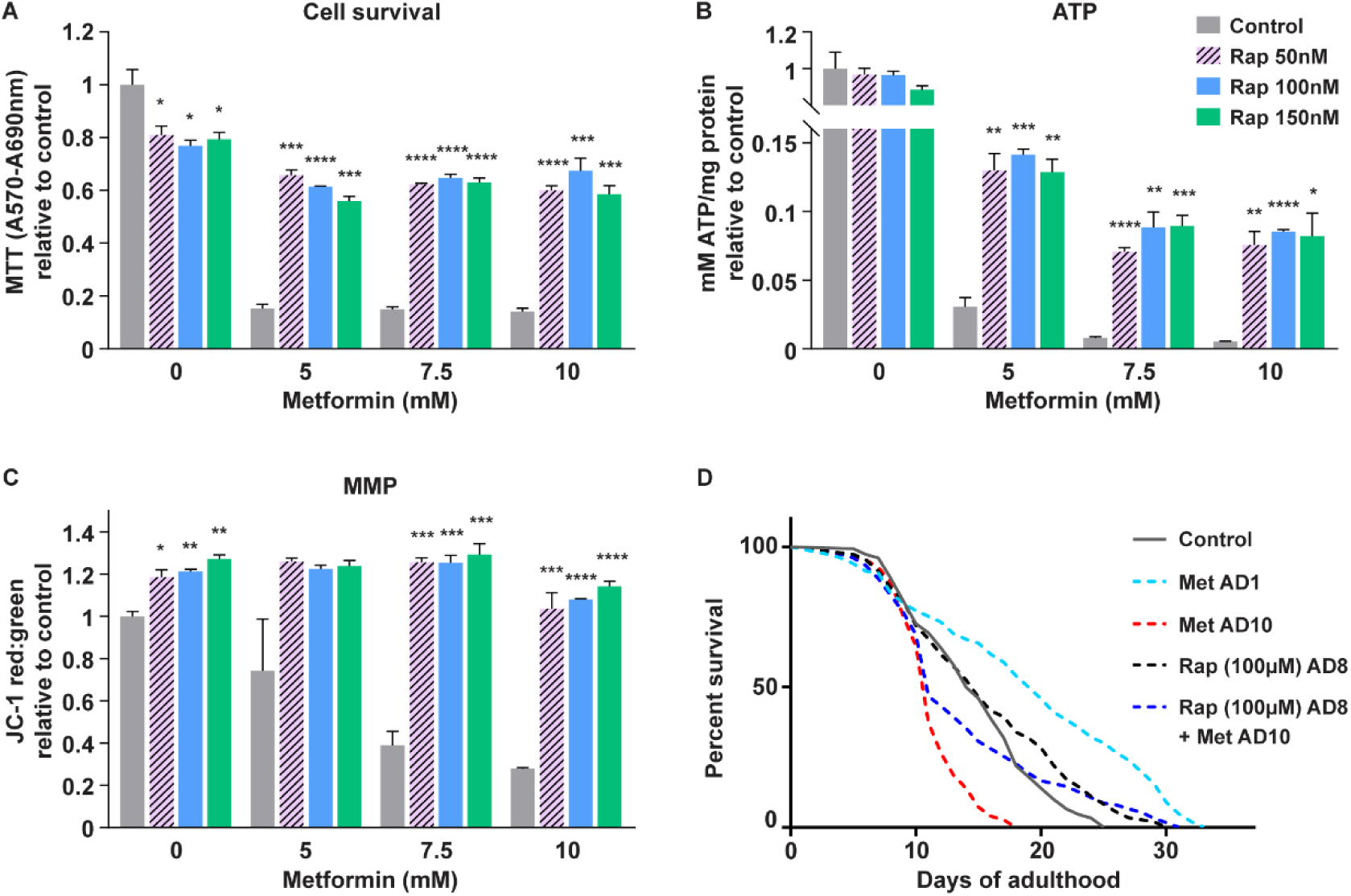
Rapamycin co-treatment alleviates ATP exhaustion and toxicity triggered by metformin in late life. Pre-senescent (PD44) primary human fibroblasts were pre-treated with indicated doses of rapamycin (Rap) for 24h followed by co-treatment with indicated doses of metformin for 24 (**A**) and 20 (**B, C**) hours; cell survival (**A**), ATP content (**B**) and mitochondrial membrane potential (**C**) were measured; values are relative to the respective untreated control (no rapamycin, no metformin) for each assay. (**D**) WT nematodes were pre-treated with 100μM rapamycin from adulthood day 8 (AD8) followed by co-treatment with 50mM metformin from AD10, survival was scored daily. For **D** significance was measured by log-rank test; all n numbers (n≥100 in all cases) and statistical values are presented in Table S1. For **A-C**, n=3 mean and SEM are presented, two-tailed unpaired t-test was used for the statistical analysis, all statistical values are presented in Table S2; * p<0,05; ** p<0,01; *** p<0,001; **** p<0,0001.

## Discussion

This study, performed in nematodes and human cells, demonstrates for the first time that metformin treatment initiated in late life triggers a range of aging-specific metabolic failures culminating in fatal ATP exhaustion (Figure S16). We show that ETC complex I inhibitor rotenone acts comparably to metformin in inducing aging-associated metabolic decline, suggesting that metformin promotes metabolic deterioration in late life via its known capacity to directly inhibit oxidative phosphorylation (Wheaton et al., 2014). The premature induction of mitochondrial impairments, comparable to aging-induced distortions of this organelle, was sufficient to trigger metformin toxicity independently of age, in line with the primary role of aging-associated mitochondrial dysfunction in late life adverse effects of metformin. The stronger negative impact of metformin on mitochondrial integrity in old cells was also indeed detected. We demonstrate that, in addition to mitochondrial alterations, the adverse effect of metformin on energy metabolism in old cells is exacerbated by the reduced capacity of these cells to upregulate glycolysis in response to metformin and by the lack of a DR mimetic lipid turnover response in old metformin exposed animals. Consistently, old but not young animals experience persistent ATP exhaustion in response to metformin which limits their cell viability and contributes to life span shortening in these animals.

By gene inactivation tests we demonstrated that AMPK mitigates the aberrant lipid accumulation in old metformin exposed animals, consistent with the known function of this kinase in restricting lipid synthesis in response to metformin (Zang et al., 2004). Strikingly, the young age dietary restriction (DR) effect of metformin was not inhibited but enhanced by the loss of AMPK gene function, contrary to the previously anticipated activator role of AMPK in this longevity-assurance response. Subsequently, we found that the DR-like effect of metformin in early life is mediated by the protein kinase A and its downstream triglyceride lipase ATGL-1, similar to the lipid turnover phenotype induced by fasting (Lee et al., 2014). Interestingly, AMPK is known to have an inhibitory effect on ATGL-1, preventing the excessive lipid exhaustion by this lipase with critical effects on longevity (Narbonne and Roy, 2009). Our data supports the model where AMPK has a similar inhibitory function in the early life DR response to metformin (Figure S16), likely explaining the well documented importance of AMPK for life-long metformin benefits. In summary, we discovered a novel previously unknown relationship between metformin and the protein kinase A pathway in mediating the beneficial DR effect of metformin. We also obtained evidence of the previously unexplored inhibitory role of AMPK in this process, which may contribute to the key protective function of AMPK in distinct metabolic stress scenarios such as metformin exposure.

Because metformin doesn’t show the toxicity comparable to our findings in elderly diabetes patients, we conducted metformin tests in old *daf-2(e1370)* mutant nematodes carrying the diabetes-like insufficiency of the *C. elegans* insulin receptor. The insulin receptor mutants indeed showed prolonged resilience to late life metformin toxicity, linked with their ability to sustain stable ATP synthesis during late life metformin exposure. This capacity is regulated by the DAF-16/FOXO stress response pathway and likely represents the adaptation of the mutants to persistent metabolic stress, prior to metformin treatment. Our data thus indicates that strong metabolic differences between non-diabetic and diabetic individuals may significantly hinder the extrapolation of metformin benefits observed in diabetes patients, including life span extension at old age, to healthy elderly humans.

Finally, we found that metformin treatment triggers a plethora of adaptive stress responses in young animals while old nematodes failed to activate these pathways to a similar extent. These data are consistent with the previously proposed key role of stress and metabolic adaptations in life extension by metformin (De Haes et al., 2014; Onken and Driscoll, 2010). The inferior induction of longevity assurance pathways, including the DR response, by metformin in old animals indicates that health extension properties of this drug may be reduced at old age in addition to metabolic failures triggered by metformin in non-diabetic old subjects. In line with the key role of early life adaptations in metformin-induced longevity extension, we found that metformin treatment restricted to early adulthood is sufficient to prolong survival (Figure S10G).

Collectively, we identified important age-specific effects of metformin which may limit therapeutic benefits of this drug for non-diabetic elderly patients. In fact, observations consistent with a reversal of metformin benefits at old age (Alfaras et al., 2017) and a negative impact of late life metformin exposure on adaptive, motor and cognitive abilities (Konopka et al., 2019; Thangthaeng et al., 2017) have recently emerged from descriptive studies performed in mice and humans. In this context, our work provides for the first time the definite molecular and genetic evidence linking age-specific adverse effects of metformin to reduced longevity assurance capacity and impaired metabolic plasticity of non-diabetic old subjects. This new knowledge is of key importance given current proposals to use metformin for improving health span of metabolic-healthy elderly humans.

## Supporting information

Proteomics data description

Lipidomics data description

## Acknowledgements

We thank Prof. KL Rudolph and Prof. Helen Morrison for critical discussions that were very helpful and important throughout the study and during manuscript preparation. We also thank the Proteomics Core Facility at FLI for supporting this study. The FLI is a member of the Leibniz Association and is financially supported by the Federal Government of Germany and the State of Thuringia. AM is supported by the German Research Council (Deutsche Forschungsgemeinschaft, DFG) via the Research Training Group Adaptive Stress Responses (GRK 1715), TP is supported by the German Academic Exchange Services (Deutsche Akademische Austauschdienst, DAAD).

## Author contribution

ME conceptualized and designed the study; LE, AD, PC, AM, TP, AN, YS performed experiments; ME, LE, AD, PC analyzed the data; ND and NR performed sample curation for proteomics analysis; JK and AO developed proteomics analysis protocols and analyzed proteomics data; AK designed the lipidomics study; LM and AK performed the lipidomics analysis; AK, OW and ME interpreted the lipidomics data; LE, PC and AO prepared figures; AD performed statistical analysis; ME wrote the manuscript and LE, AD, PC, JK, AK and AO reviewed the manuscript.

## Data availability

The mass spectrometry proteomics data that support the findings of this study have been deposited to the ProteomeXchange with the data set identifier PXD011579. All other relevant data is available from the corresponding author.

## Declaration of interests

Authors declare no competing interests.

## Supplemental Materials

### Materials and methods

#### Worm strains

All strains were cultured according to standard conditions. Strains used were N2 Bristol (wild type), RB754 *aak-2(ok524)* X, VC3201 *atfs-1(gk3094)* V, QV225 *skn-1(zj15)* IV, MQ887 *isp-1(qm150)* IV, CB1370 *daf-2(e1370)* III and CB4037 *glp-1(e2141)* III. The following additional strains were used in the supplementary figures VC2654 *ubl-5(ok3389)* I, SJ4100 *zcls13 [hsp-6::GFP]* V, MAE10 *daf-2(e1370);daf-16(mu86)*, TJ356 *zIs356 [daf-16p::daf-16a/b::GFP + rol-6(su1006)]* IV and VK1093 *vkEx1093 [nhx-2p::mCherry::lgg-1]*.

#### Bacterial strains and growth conditions

*E. coli* OP50 and *E. coli* HT115(DE3) were seeded on lysogeny broth (LB) agar plates and kept at 4 °C. Overnight cultures were grown at 37 °C in LB media without antibiotics and provided to worms on standard nematode growth medium (NGM) agar. The supplements were given in NGM agar: metformin (M2009 TCI), FCCP (15218 Cayman Chemical Company) and rapamycin (R0395 Sigma, R-5000 LC Laboratories). UV-killing was performed 24h after bacteria seeding by exposing the surface of the plates, without lids, to UV for 5 minutes (ChemiDOC XRS+ Bio-Rad); loss of viability was confirmed by subsequent liquid culture.

#### RNA interference treatments

HT115 bacteria containing specific RNAi constructs were grown on lysogeny broth agar plates supplemented with ampicillin and tetracycline (Roth). Plates were kept at 4 °C. Overnight cultures were grown in lysogeny broth media containing ampicillin. RNAi expression was induced by adding 1 mM isopropylthiogalactoside (IPTG, Sigma) and incubating the cultures at 37 °C for 20 min before seeding bacteria on NGM agar supplemented with ampicillin and 3 mM IPTG.

#### Lifespan analysis

The experiments were carried out at 20°C under standard laboratory conditions, unless stated otherwise. Synchronized L4 larvae were placed on 60 mm dishes containing NGM agar and bacteria at a density of at least 70 worms per plate, two plates per condition (n≥140). Worms were transferred to new plates every other day until adulthood day 7 (AD7) and later transferred to new plates every 3 days. The number of dead animals was scored daily. Treatment with supplements including metformin was initiated by transfer of animals onto plates containing respective compounds. The analysis of the lifespan data including statistics was performed by using GraphPad Prism 7 software.

#### Sample preparation for proteomic analysis

Approximately 500 age-synchronized worms per sample were used. Worms were collected in M9 buffer and pelleted; pellets were frozen at −80°C. Upon thawing the sample volume was adjusted to 100 µL with PBS; samples were suspended in 2x lysis buffer (2x = 0.2M HEPES/pH 8; 2% SDC; 0.2M DTT; 2mM EDTA) and subjected to a freeze-thaw cycle in liquid nitrogen followed by incubation at 95 °C with shaking at 500 rpm for 5 minutes. Samples were then sonicated in a Bioruptor (Diagenode, Beligum) (10 cycles with 1 minute on and 30s off with high intensity at 20°C). The lysates were clarified, and debris precipitated by centrifugation at 14000 rpm for 10 minutes. For reduction and alkylation of the proteins, the lysate supernatants were first incubated at 56 °C for 30 min with DTT (5 µL of a 100 mM solution) and subsequently treated with iodacetamide (room temperature, in the dark, 20 minutes, 5 µL of a 200 mM solution). 10 % of the sample was removed to check lysis on a coomassie gel. The remaining 90% of the samples was treated with 1 volume ice cold 100% TCA to 4 volumes sample and left to stand on ice for 30 minutes to precipitate the proteins. The samples were then centrifuged at 14000 rpm for 20 minutes, 4 °C. After removal of the supernatant, the precipitates were washed once with 1000 µL 10% TCA, vortexed, centrifuged again for 20 minutes at 4°C then washed twice (2 x 1000 µL) with ice cold (stored at −20 °C before use) acetone. Vortexing and centrifugation steps repeated as before. The pellets were then allowed to air-dry before being dissolved in digestion buffer (50 µL, 3M urea in 0.1M HEPES, pH 8; 1 µg LysC) and incubated for 4 h at 37 °C. Then the samples were diluted 1:1 with Milli-Q water (to reach 1.5 M urea) and were incubated with 1 µg trypsin for 16 h at 37 °C. The digests were then acidified with 10% trifluoroacetic acid and then desalted with Waters Oasis® HLB µElution Plate 30 µm (Waters Corporation, Milford, MA, USA) in the presence of a slow vacuum. In this process, the columns were conditioned with 3×100 µL solvent B (80% acetonitrile; 0.05% formic acid) and equilibrated with 3x 100 µL solvent A (0.05% formic acid in Milli-Q water). The samples were loaded, washed 3 times with 100 µL solvent A, and then eluted into PCR tubes with 50 µL solvent B. The eluates were dried down with the speed vacuum centrifuge and dissolved in 50 µL 5% acetonitrile, 95% Milli-Q water, with 0.1% formic acid. 20 µL was transferred to an MS vial and 1 µL of HRM kit peptides (Biognosys, Zurich, Switzerland) was spiked into each sample prior to analysis by LC-MS/MS.

#### LC-MS/MS

Peptides were separated using the nanoAcquity UPLC system (Waters) fitted with a trapping (nanoAcquity Symmetry C_18_, 5µm, 180 µm x 20 mm) and an analytical column (nanoAcquity BEH C_18_, 1.7µm, 75µm x 250mm). The outlet of the analytical column was coupled directly to an Orbitrap Fusion Lumos (Thermo Fisher Scientific) using the Proxeon nanospray source. Solvent A was water, 0.1 % formic acid and solvent B was acetonitrile, 0.1 % formic acid. The samples (approx. 1 µg for DDA and 3 µg for DIA) were loaded with a constant flow of solvent A at 5 µL/min onto the trapping column. Trapping time was 6 minutes. Peptides were eluted via the analytical column with a constant flow of 0.3 µL/min. During the elution step, the percentage of solvent B increased in a non-linear fashion from 0 % to 40 % in 120 minutes. Total runtime was 145 minutes, including clean-up and column re-equilibration. The peptides were introduced into the mass spectrometer via a Pico-Tip Emitter 360 µm OD x 20 µm ID; 10 µm tip (New Objective) and a spray voltage of 2.2 kV was applied. The capillary temperature was set at 300 °C. The RF lens was set to 30%. Data from each sample were first acquired in DDA mode for one of the replicates in order to create a spectral library. The conditions were as follows: Full scan MS spectra with mass range 350-1650 *m/z* were acquired in profile mode in the Orbitrap with resolution of 60000. The filling time was set at maximum of 50 ms with limitation of 2e5 ions. The “Top Speed” method was employed to take the maximum number of precursor ions (with an intensity threshold of 5e4) from the full scan MS for fragmentation (using HCD collision energy, 30%) and quadrupole isolation (1.4 Da window) and measurement in the Orbitrap (resolution 15000, fixed first mass 120 *m/z*), with a cycle time of 3 seconds. The MIPS (monoisotopic precursor selection) peptide algorithm was employed but with relaxed restrictions when too few precursors meeting the criteria were found. The fragmentation was performed after accumulation of 2e5 ions or after filling time of 22 ms for each precursor ion (whichever occurred first). MS/MS data were acquired in centroid mode. Only multiply charged (2^+^ - 7^+^) precursor ions were selected for MS/MS. Dynamic exclusion was employed with maximum retention period of 15 s and relative mass window of 10 ppm. Isotopes were excluded. In order to improve the mass accuracy, internal lock mass correction using a background ion (*m/z* 445.12003) was applied. For data acquisition and processing of the raw data Xcalibur 4.0 (Thermo Scientific) and Tune version 2.1 were employed.

For the DIA data acquisition the same gradient conditions were applied to the LC as for the DDA and the MS conditions were varied as described: Full scan MS spectra with mass range 350-1650 *m/z* were acquired in profile mode in the Orbitrap with resolution of 120000. The filling time was set at maximum of 20 ms with limitation of 5e5 ions. DIA scans were acquired with 34 mass window segments of differing widths across the MS1 mass range with a cycle time of 3 seconds. HCD fragmentation (30% collision energy) was applied and MS/MS spectra were acquired in the Orbitrap with a resolution of 30000 over the mass range 200-2000 *m/z* after accumulation of 2e510^5^ ions or after filling time of 70 ms (whichever occurred first). Ions were injected for all available parallelizable time). Data were acquired in profile mode.

For the second experiment (young wild-type, *daf-2(e1370)* and *daf-2(e1370); daf-16(mu86)* worms), the following changes were made to the data acquisition: data were acquired instead on a QE-HFX MS (Thermo), connected to an M-Class nanoacquity (Waters). Columns/solvents/gradients and source settings were all the same as for the Lumos. DDA data were acquired on a subset of samples from all conditions, with the following settings changes compared to Lumos: MS1 fill time was 20 ms, AGC target 3e6. Top N was used (=15) and the intensity threshold was 4e4. Normalized Collision Energy (NCE) with HCD was set to 27% and a 1.6 Da window was used for quadrupole isolation. MS2 data were acquired in profile mode from 200 – 2000 *m/z*. MS2 fill time was 25 ms or an AGC target of 2e5. Only 2-5+ charge states were selected for MS/MS. Dynamic exclusion was 20s, and the peptide match “preferred” option was selected. For the DIA data, LC conditions remained unchanged and the following changes were made for the MS: fill time for MS1 was 60ms, with an AGC target of 3e6. DIA segments in MS2 were acquired for 40ms or AGC target of 3e6 at a fixed first mass of 200 *m/z*. Stepped HCD NCEs of 25.5, 27 and 30 % were employed.

#### Proteomics data analysis

For library creation, the DDA data was searched using MaxQuant (version 1.5.3.28; Martinsreid, Germany). The data were searched against a species specific (*C. elegans*) UniProt database with a list of common contaminants appended, as well as the HRM peptide sequences. The data were searched with the following modifications: Carbamidomethyl (C) (Fixed) and Oxidation (M)/ Acetyl (Protein N-term) (Variable). The mass error tolerance for the full scan MS and MS/MS spectra was set at 20 ppm. A maximum of 1 missed cleavage was allowed. The identifications were filtered to satisfy a false discovery rate of 1 % on peptide and protein level.

A spectral library was created from the MaxQuant output of the DDA runs combined using Spectronaut (version 10, Biognosys, Switzerland). This library contained 58296 precursors, corresponding to 4624 protein groups using Spectronaut protein inference. DIA data were then uploaded and searched against this spectral library. For the second experiment, DDA and DIA data were directly searched in Spectronaut Pulsar X, with the same database and settings applied as for MaxQuant to create a DpD library against which this data was searched. This library contained 75365 precursors corresponding to 5287 protein groups.

Relative quantification was performed in Spectronaut for each pairwise comparison (such as young 24h metformin treated versus young 24h untreated or old 48h metformin treated versus old 48h untreated) using the triplicate samples (4-5 replicates for the second experiment) from each condition. The data (candidate table with Log2 fold changes and Q values for each pairwise comparison) was then exported to Excel (presented as Table S3) and further data analyses and visualization (such as Venn diagrams, scatter plots and box plots) were performed with R-studio (version 0.99.902) using in-house pipelines and scripts.

#### Lipidomics analysis by UPLC-MS/MS

Young and old wild type animals (n≥700) were treated with 50mM metformin (or vehicle) for 24 hours. For the analysis lipids were extracted by successive addition of PBS pH 7.4, methanol, chloroform, and saline and then separated on an Acquity UPLC BEH C8 column (1.7 μm, 2.1 × 100 mm) using an Acquity UPLC system (Waters). The UPLC system was coupled to a QTRAP 5500 mass spectrometer (Sciex) equipped with an electrospray ionization source to detect the fatty acid anion fragments of glycerophospholipids by multiple reactions monitoring in the negative ion mode. Free fatty acids were quantified by single ion monitoring in the negative ion mode, and triacylglycerols were analyzed based on transitions of [M+NH_4_]^+^ to [M-fatty acid anion]^+^ fragments. Specifically, phospholipids were analyzed as described in (Koeberle et al., 2013) and triglycerides were assessed as described in (Koeberle et al., 2015). Absolute intensities are defined as summarized signal intensities of all detected species of a given lipid type (e.g. all free fatty acids) normalized to the internal standard and the number of worms. Relative intensities are defined as proportions of individual lipid signals (e.g. individual triglyceride subtypes) relative to total signal intensity for a given lipid type (e.g. total triglycerides).

#### Lipid staining

Oil Red O (Sigma) preparation and staining was performed as previously described (Han et al., 2017). After staining the animals were mounted onto 2% agar pads and imaged by using the Axio Scan.Z1 machine (Carl Zeiss) at 20x magnification coupled to the HV-F202SCL color camera (Hitachi). The same exposure and z-stack settings were used across all conditions and experiments. The images were split by using the Zen2.6 software (Carl Zeiss) and quantified by using ImageJ (NIH): each raw image was background-subtracted, greyscale-converted and inverted for quantification as previously described (Han et al., 2017), frontal anterior areas of the intestine (half the animal length) were outlined and quantified in each case. Measurement of Oil Red O mean intensity per worm was performed using the automatic “Analyze Measure” feature of the ImageJ software. For each image (animal) the background was subtracted, and mean grey values were plotted as arbitrary units (a.u.).

#### Fluorescence microscopy

*hsp-6::GFP* worms were age-synchronized and L1 larvae were fed with bacteria containing empty vector (EV), *mip-1* RNAi (Ahringer library) or EV supplemented with 50mM metformin. Worms were incubated at 20°C and imaged on adulthood day 1 (AD1) using AxioZom_V16 microscope (Zeiss) at 90X magnification. Worms were immobilized by exposure to ice for 20 minutes prior to imaging. *daf16a/b::GFP* reporter worms were synchronized and fed with OP5O *E. coli*. On AD1, worms were transferred onto plates supplemented with 50mM metformin or control plates. Heat stress was used as a positive control: AD1 worms were exposed to 35.7°C for 3h (BF115 incubator, Binder) and imaged at indicated times at 20°C. For the old age group, worms were transferred onto metformin or control plates on AD8 and scored on AD10. We used AxioZoom_V16 microscope (Zeiss) for scoring and imaging (250X magnification), n=100 worms were imaged per each condition. *mCherry::lgg-1* reporter worms were treated with 50mM metformin or transferred to control plates and the number of puncta per animal was scored after 6, 12 and 24h of treatment. AxioZoom_V16 microscope (Zeiss) was used for imaging (160X magnification), n=10 worms were imaged per each condition.

#### ATP content

Systemic ATP levels in *C. elegans* were measured as previously described (Artal-Sanz and Tavernarakis, 2009), with small modifications. Briefly, 50 worms per sample were collected, washed with S-basal buffer, and snap frozen in 50µl S-basal buffer using liquid nitrogen before final storage at −80°C. Two biological replicates were collected per condition. AD1 and AD10 frozen worms were boiled for 15 minutes at 100°C, cooled to RT, sonicated and centrifuged to pellet debris. The supernatant was then collected to fresh tubes and diluted tenfold before measurement. 50µl of each sample was transferred in triplicate to a 96 well flat bottom black plate and ATP content was determined using Roche ATP Bioluminescence Assay Kit HSII (Sigma) and a luminometer plate reader with injection system (Mithras LB940, Berthold Technologies). 50µl of luciferase reagent was injected per well and the light signal was integrated for 10s after a delay of 1s. ATP levels were normalized to total protein content (A280nm in NanoDrop (Thermo Fischer). Extraction of ATP from human cells was performed according to the kit supplier’s protocol. Briefly, cells were seeded and treated in 96 well plates and, after treatment, culture media was removed and cells were washed twice with HBSS to remove any traces of FBS and lysed with 100 µl of cell lysis reagent (Roche ATP Bioluminescence Assay Kit HSII, Sigma). Lysate supernatant was diluted tenfold and measurement was done as explained above for worms. Resulting ATP levels were normalized to total protein content determined with Pierce BCA Protein Assay Kit (Thermo Fischer).

#### Cell culture

BJ human skin fibroblasts were maintained in DMEM high glucose media with L-glutamine and pyruvate supplemented with 10% of fetal bovine serum (FBS) (all from Sigma). For cell culture assays cells were seeded in complete DMEM on 96 well plates, flat bottom, at a density of 20.000 cells per well. Cells were left overnight for proper cell attachment and the different treatments were then performed for the times indicated. For pre-treatment with rapamycin complete DMEM was used. For treatment with metformin (plus or without supplements), cells were first washed with HBSS and media was replaced with DMEM without glucose, L-glutamine or phenol red, and supplemented with 10% FBS. For FCCP (Sigma) and rapamycin co-treatments, control cells were treated with vehicle dimethylsulfoxide (DMSO, Sigma). DMSO concentrations did not exceed 0,05%. All incubations and culturing were done in a humidified 37°C, 5% CO2 incubator (BBD6220, Thermo Scientific).

#### Cell survival

25µl of a filtered 5mg/mL stock solution of MTT (Sigma) was added to each cell containing well to a final concentration of 1mg/mL. Plates were wrapped in aluminum foil to protect from light and incubated for 2h (37 °C, 5% CO2). MTT media was then removed and replaced by MTT solubilization solution (0.1N HCL in absolute isopropanol). Absorbance of the converted dye was measured at 570nm with background subtraction at 690nm using a plate-reading spectrophotometer (Infinite M200 Pro, Tecan).

#### Cell death

Quantification of cell death was performed with the Pierce LDH Cytotoxicity Assay kit (Thermo Scientific). After cell treatment 50µl of each sample medium was transferred to a new 96 well plate (flat bottom) and mixed with 50µl of Reaction Mixture. After 30-minute incubation at RT 50µl of Stop Solution was added. Absorbance was measured at 490nm with background subtraction at 680nm using a plate-reading spectrophotometer (Infinite M200 Pro, Tecan). % of cytotoxicity was calculated as indicated in the kit.

#### Mitochondrial membrane potential

Drug-induced changes of mitochondrial membrane potential (MMP) were assessed using JC-1 dye (Thermo Fischer). In cells with healthy mitochondria, JC-1 forms J-aggregates with red fluorescence. However, in damaged cells with low MMP JC-1 remains in the green fluorescent monomeric form. Fluorescence intensities were measured with a plate-reading spectrophotometer (Infinite M200 Pro, Tecan) and loss of MMP was determined by calculating JC-1 red:green fluorescence ratio.

#### Seahorse analysis

Mitochondria function parameters such as the oxygen consumption rate (OCR) and extracellular acidification rate (ECAR) were measured in real time, using Agilent Seahorse XF24 Analyzer according to the protocol provided by Agilent Technologies. Briefly, cells were seeded in XF24 TC-treated cell culture microplates (Agilent Technologies) and left overnight to adhere and grow until confluence. As for the other cell assays in this manuscript, cells were seeded in complete DMEM and further treated with metformin in DMEM with 10%FBS but without glucose, L-glutamine or phenol red for the indicated time. After treatment, cells were washed 2x and incubated for 1h at 37°C in a non-CO2 incubator with XF base media (Agilent Technologies) supplemented with glutamine 2mM, pyruvate 1mM and glucose 10mM (all Agilent Technologies). The sensor cartridge of the XFe24 extracellular flux assay kit (Agilent Technologies) was left hydrating in XF calibrant at 37°C in a non-CO2 incubator overnight. The modulators of respiration were prepared fresh and loaded into the sensor cartridge injection ports immediately before the assay at the following concentrations: Oligomycin 1µM (port 1), FCCP 6 µM (port2) and Rotenone/Antimycin-A 0,5 µM each (port 3) (all from Sigma). Assays were run and result analyses were done using Seahorse Wave software (Agilent Technologies). Equal cell number in young and old cultures was ensured by nuclear (DAPI, Sigma) staining prior to running the tests. All calculated values were normalized to total protein content of exact corresponding wells measured with Pierce BCA Protein Assay Kit (Thermo Fischer).

#### Statistical analysis

For all lifespan experiments, median survival was calculated and statistical significance was determined by using both log-rank (Mantel-Cox) and Gehan-Breslow-Wilcoxon tests. For lipid content, ATP levels, survival, cell death and JC-1 assays unpaired t-tests with two-tailed distribution were performed to evaluate statistical significance. Error bars represent standard deviation of the mean in all cases. Statistical calculations were performed by using GraphPad Prism 7 in all cases except proteomics analysis. Worms of a given genotype were randomly selected from large populations for each experiment without any preconditioning. Blinding was not applied as the experiments were carried out under highly standardized and predefined conditions such that an investigator-induced bias can be excluded.

**Figure S1.**
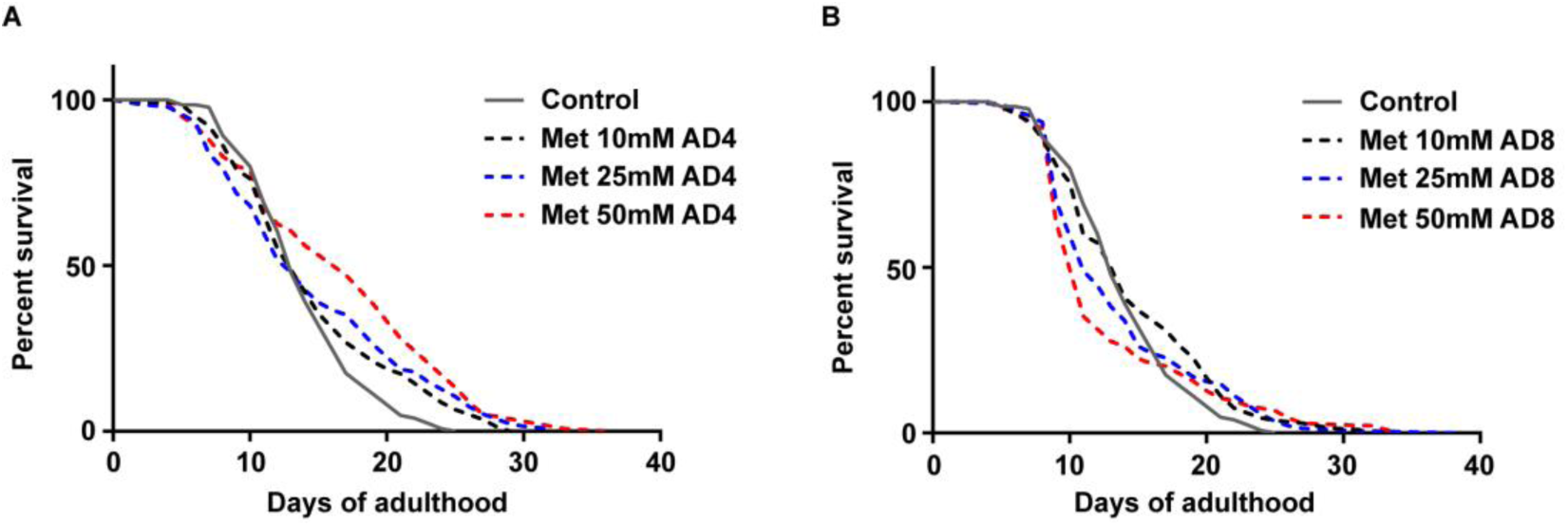
Life extending properties of metformin deteriorate with age. Wild type (WT, N2 Bristol strain) worms were treated with indicated doses of metformin (Met) from day 4 (**A**) and day 8 (**B**) of adulthood (AD4 and AD8 respectively), survival was scored daily. Significance was determined by log-rank test, all n numbers (n≥100 in all cases) and statistical values are presented in Table S1. Each experiment was repeated ≥3 times; one representative result is shown in each case.

**Figure S2.**
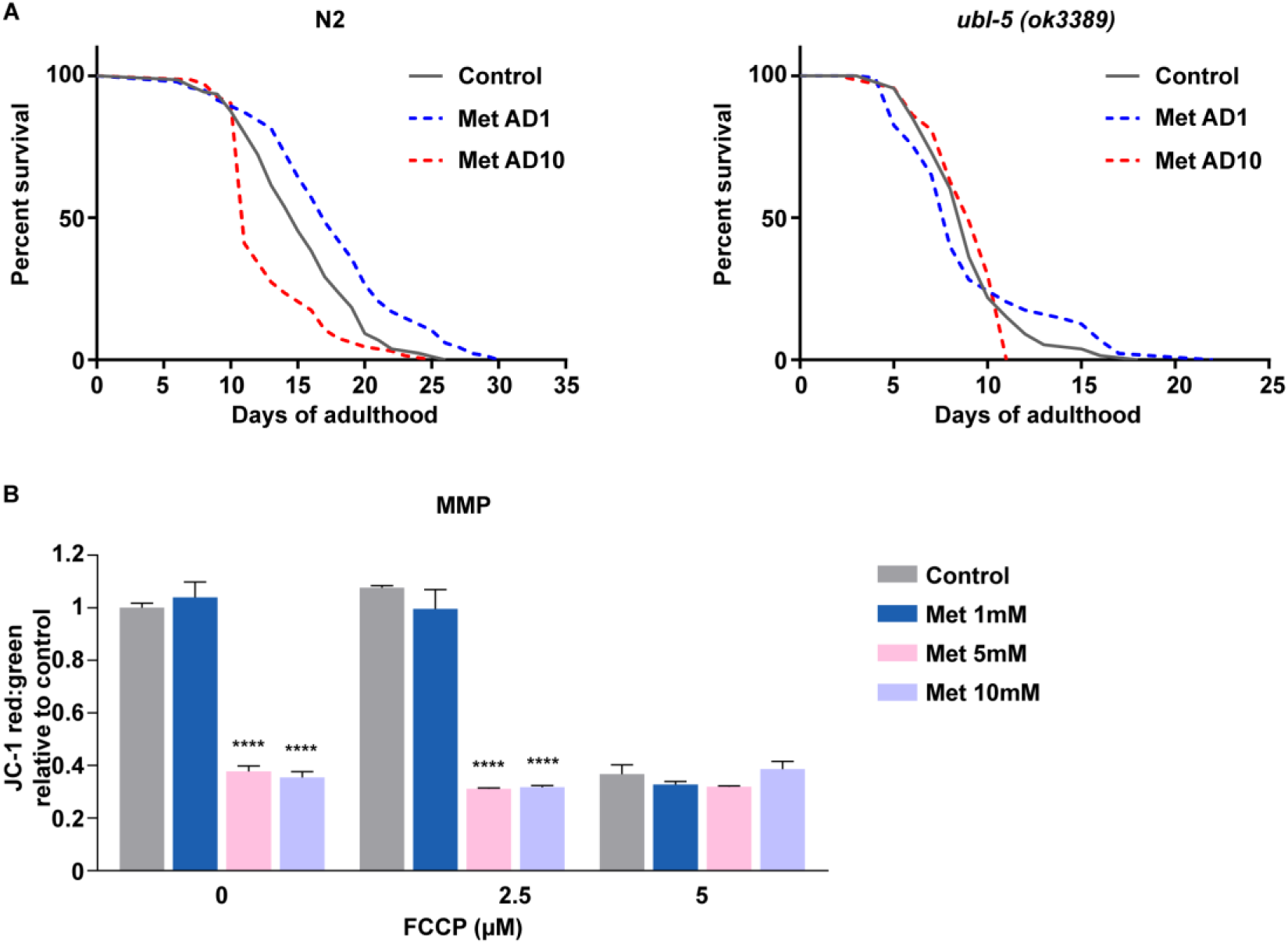
Mitochondrial impairments pre-dispose nematodes and human cells to metformin toxicity. (**A**) WT animals (**left panel**) and *ubl-5(ok3389)* mutants (**right panel**) were treated with 50mM metformin on days 1 and 10 of adulthood (AD1 and AD10 respectively), survival was scored daily. (**B**) Early passage (population doubling, PD37) fibroblasts were treated with indicated doses of metformin (Met) in absence or presence of indicated concentrations of FCCP for 24 hours; mitochondrial membrane potential was measured by JC-1 staining. All values are relative to the untreated control (no FCCP, no metformin). DMSO was used as vehicle control for FCCP, the own effect of the vehicle is seen at 0μM FCCP. Grey bars depict the effect of FCCP alone (metformin-independent) on the mitochondrial membrane potential (MMP), proof of concept MMP decline is seen at 5μM FCCP. For **A** significance was calculated by log-rank test, all n numbers (n≥100 in all cases) and statistical values are presented in Table S1; for **B** n=3, mean and SEM are presented; two-tailed unpaired t-test was used for the statistical analysis, statistical values are shown in Table S2; **** p<0,0001.

**Figure S3.**
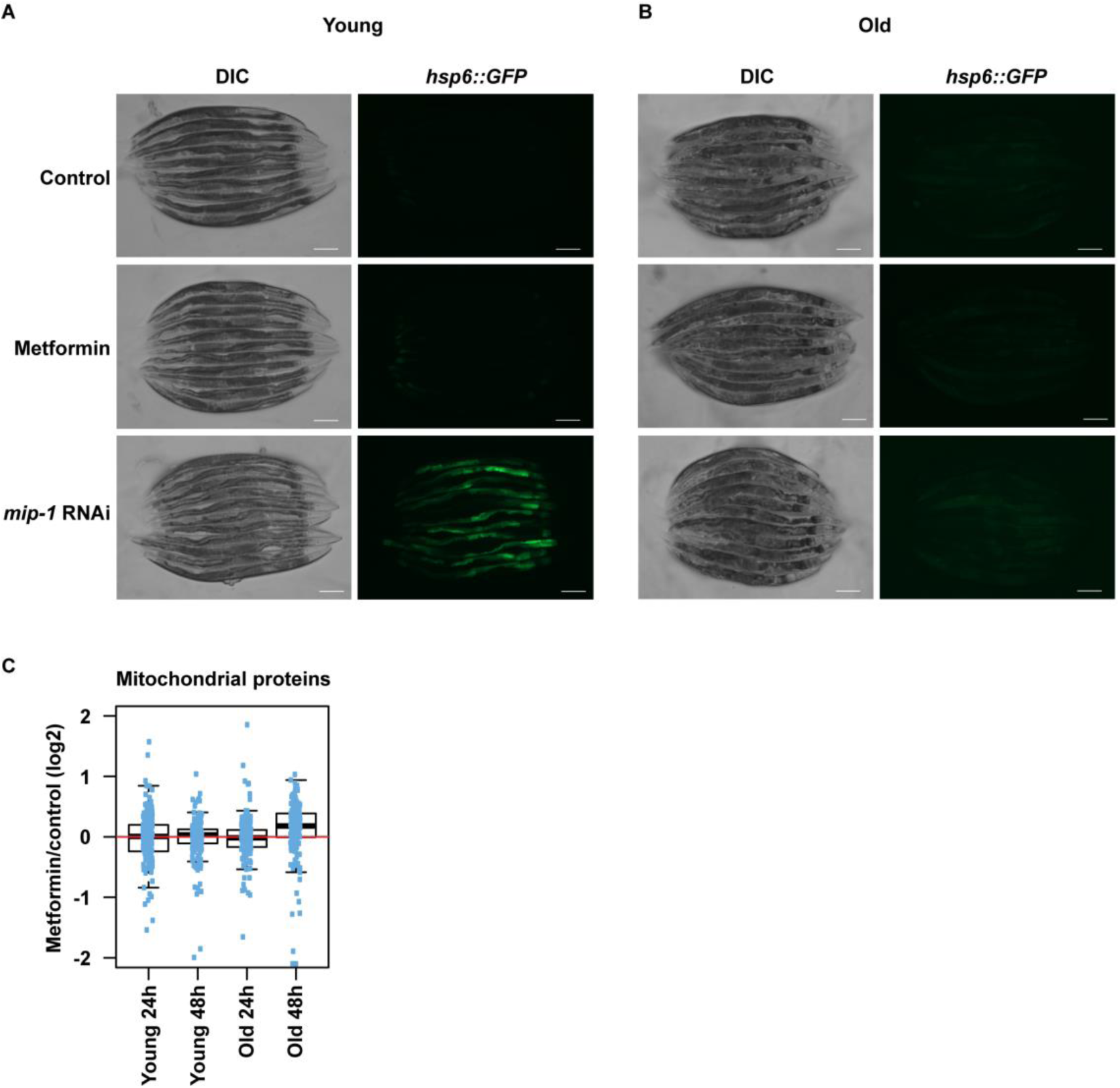
Metformin treatment doesn’t induce mitochondrial unfolded protein response or a decrease of mitochondrial content indicative of mitophagy. Day 1 (**A**) and day 10 (**B**) adult transgenic nematodes (young and old respectively) expressing GFP under control of the *hsp-6* gene promoter were treated with 50mM metformin or grown in presence of *mip-1* RNAi (positive control). GFP expression was assessed by microscopy after 24h of metformin treatment. The experiment was repeated 3 times, representative result is shown, scale bar is 200µm. (**C**) Young and old animals were treated with 50mM metformin and changes of mitochondrial protein levels were measured by mass spectrometry at 24h and 48h of treatment. Dots represent individual mitochondrial proteins, mean fold change value (taking all detected mitochondrial proteins in consideration) is shown in bold, the upper and lower limits of the boxplot indicate the first and third quartile, respectively, and whiskers extend 1.5 times the interquartile range from the limits of the box. All fold changes were calculated against age- and time point matched untreated controls (for both 24h and 48h of treatment). The complete list of proteins used for the boxplot analysis along with individual fold changes and statistical values is reported in Table S3.

**Figure S4.**
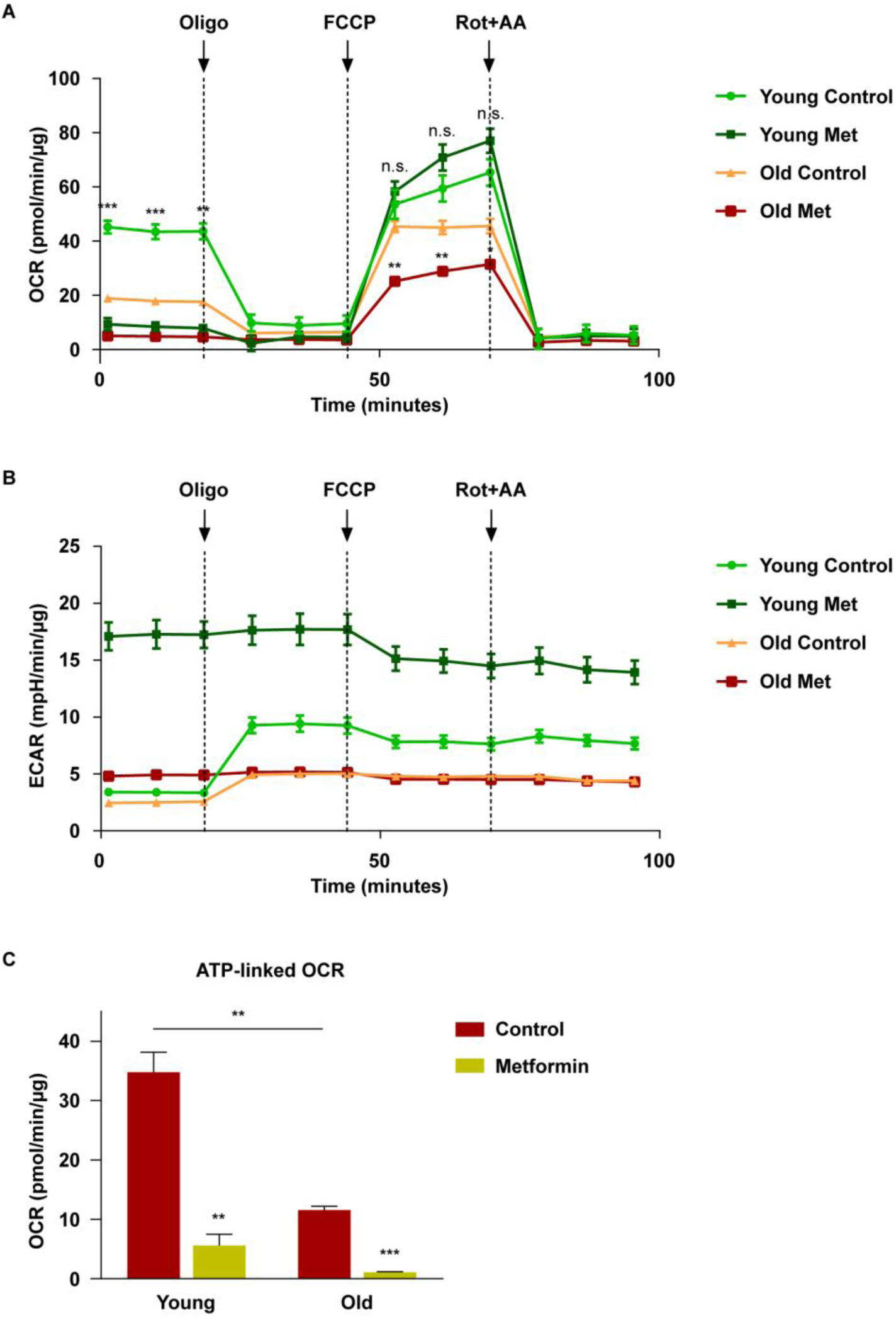
Metformin induces a distortion of energy metabolism in old human primary cells. Young (PD29) and old (PD61) primary human skin fibroblasts were treated with metformin for 16h, then washed and pre-incubated with media containing high glucose (10mM), glutamine (2mM) and pyruvate (1mM); Seahorse analysis of OXPHOS (**A**) and glycolysis (**B**) was performed by measuring continuous oxygen consumption rates (OCR) and extracellular acidification rates (ECAR), respectively, following injection of oligomycin (1mM), FCCP (3mM), antimycin-A (0,5mM) and rotenone (0,5mM). The recorded values were normalized to protein content. (**C**) ATP-linked OCR was calculated by comparison of basal OCR and OCR following oligomycin (ATP synthase inhibitor) injection. For **A-C** n=3, mean and SEM are presented; two-tailed unpaired t-test was used for the statistical analysis (**A**, **C**), all statistical values are presented in Table S2; * p<0,05; ** p<0,01; *** p<0,001.

**Figure S5.**
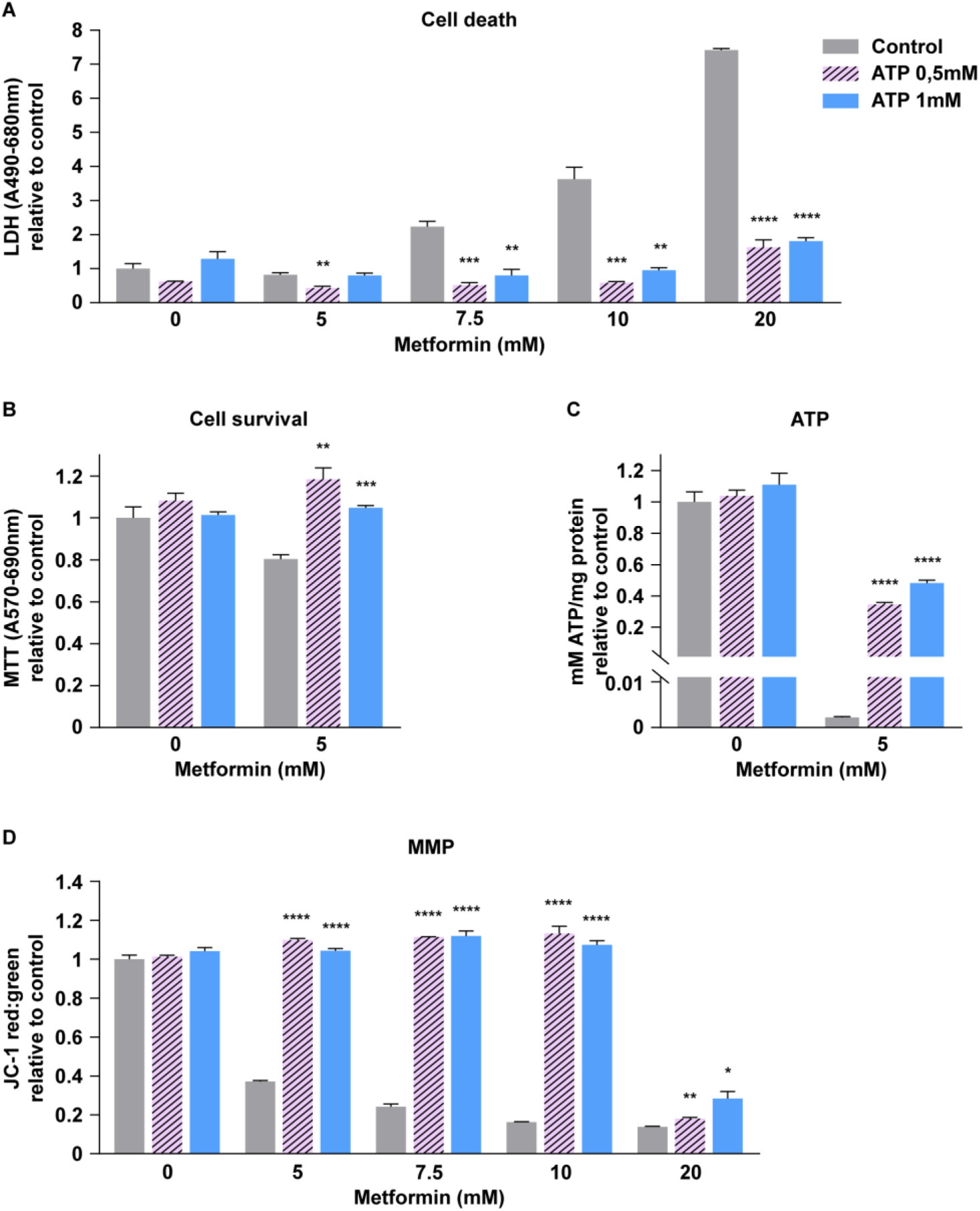
Ectopic ATP supplementation alleviates metformin toxicity in human primary fibroblasts. Pre-senescent (PD44) primary human skin fibroblasts were treated with indicated doses of metformin in presence or absence of indicated concentrations of ATP for 24h hours; cell death (**A**, LDH assay), cell survival (**B**, MTT assay), ATP content (**C**) and mitochondrial membrane potential (**D**, JC-1 assay) were measured; the data are complementary to Fig. 3I-J; values are relative to the respective untreated control (no ATP, no metformin) for each assay. The data from cells treated with 5mM metformin demonstrates that ATP exhaustion and loss of mitochondrial membrane potential precede metformin-triggered cell death. n=3, mean and SEM are presented, two-tailed unpaired t-test was used for the statistical analysis, statistical values are shown in Table S2; * p<0,05; ** p<0,01; *** p<0,001; **** p<0,0001.

**Figure S6.**
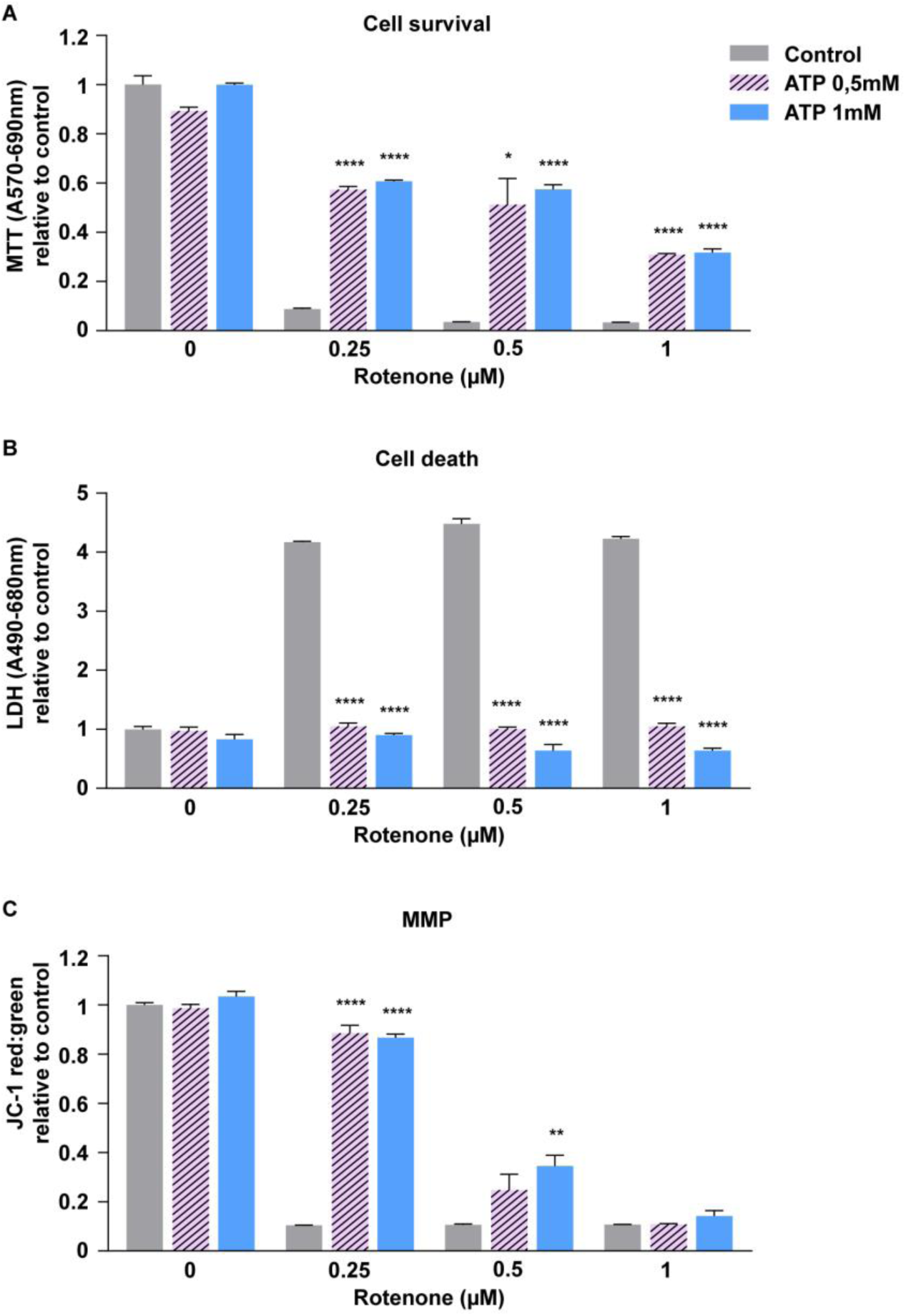
Ectopic ATP supplementation alleviates the toxicity of ETC complex I inhibitor rotenone in human primary fibroblasts. Pre-senescent primary human skin fibroblasts were treated with indicated doses of rotenone for 21 hours; cell survival (**A**, MTT assay), cell death (**B**, LDH assay), and mitochondrial membrane potential (**C**, JC-1 assay) were measured. Values relative to the respective untreated control (no ATP, no rotenone) are shown for all assays. n=3, mean and SEM are presented, two-tailed unpaired t-test was used for the statistical analysis, statistical values are shown in Table S2; * p<0,05; ** p<0,01; **** p<0,0001.

**Figure S7.**
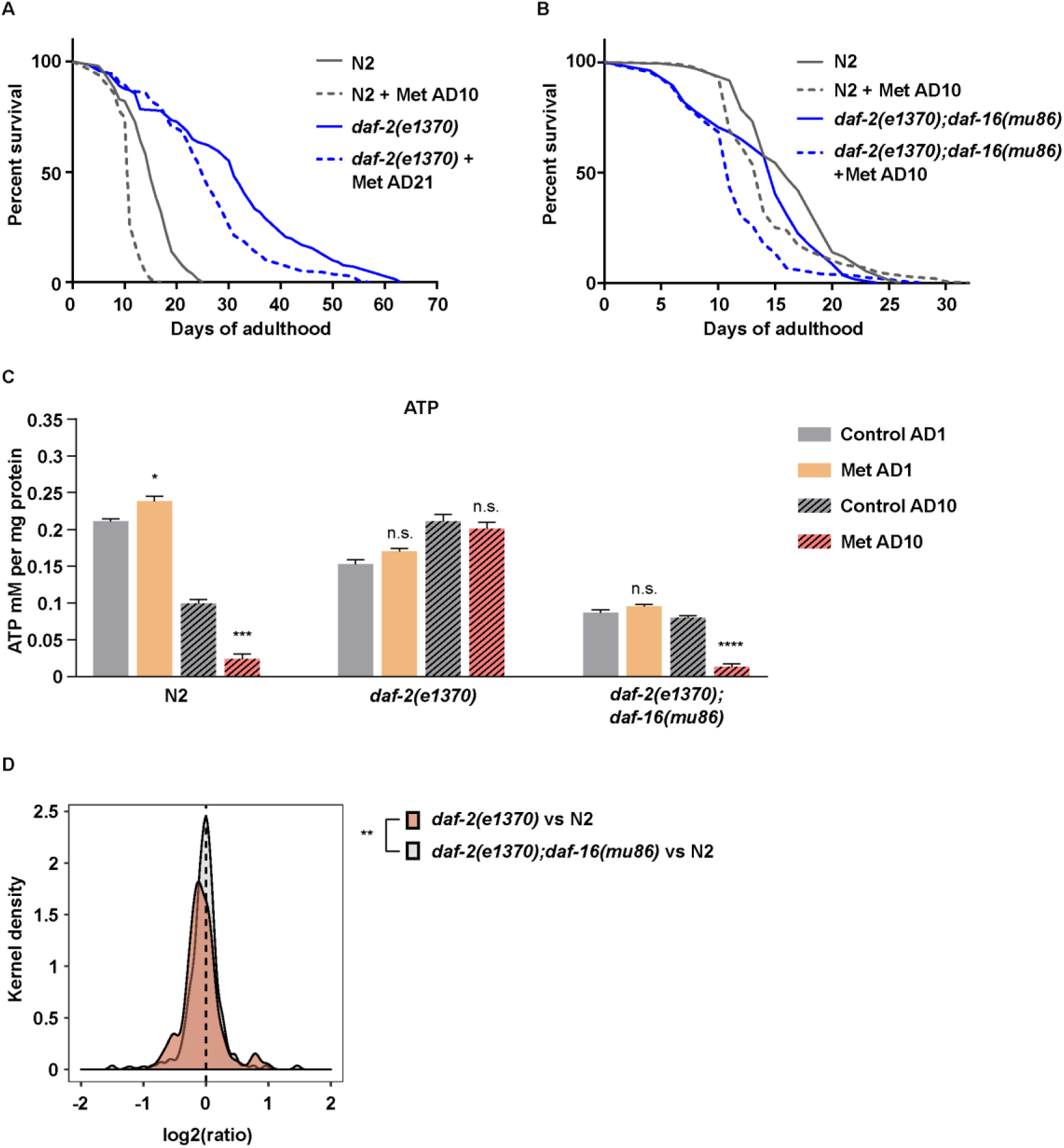
DAF-16/FOXO is required for the enhanced resistance of insulin receptor deficient animals to late life metformin toxicity. (**A**) *daf-2(e1370)* mutant animals were treated with 50mM metformin (Met) on adulthood day 21 (AD21) along with AD10 wild type (N2 Bristol strain) control animals, survival was scored daily. (**B**) *daf-2(e1370)*;*daf-16(mu86)* mutant animals were treated with 50mM metformin on adulthood day 10 (AD10) along with age-matched wild type controls, survival was scored daily. (**C**) Whole organism ATP levels were measured in wild type, *daf-2(e1370)* and *daf-2(e1370)*;*daf-16(mu86)* mutant animals treated with 50mM metformin on AD1 and AD10, the measurement was performed after 36h of metformin exposure; the data are complementary to Fig. 4C, all presented whole animal ATP measurements were performed in parallel to ensure comparability. (**D**) The proteome of young (AD2) wild type, *daf-2(e1370)* and *daf-2(e1370)*;*daf-16(mu86)* animals was analyzed by mass spectrometry; log_2_ fold changes (relative to wild type) of mitochondrial proteins (214 in total) were compared between the *daf-2(e1370)* and *daf-2(e1370)*;*daf-16(mu86)* mutant strains. The distribution of log_2_ fold changes shows reduced abundance of mitochondrial proteins in *daf-2(e1370)* mutants, while *daf-2(e1370)*;*daf-16(mu86)* mutants show mitochondrial protein levels comparable to wild type. For **A**-**B** significance was measured by log-rank test; n numbers (n≥100 in all cases) and statistical values are presented in Table S1. For **C** n=100, mean and SEM are presented, two-tailed unpaired t-test was used for the statistical analysis, all statistical values are presented in Table S2. For **D** 5 independent pools of worms (n≥500 in each) were measured for each strain, the complete data is presented in Table S3. The Wilcoxon rank sum test was used to assess significance between the log_2_ fold change distributions. * p<0.05; *** p<0.001; **** p<0.0001.

**Figure S8.**
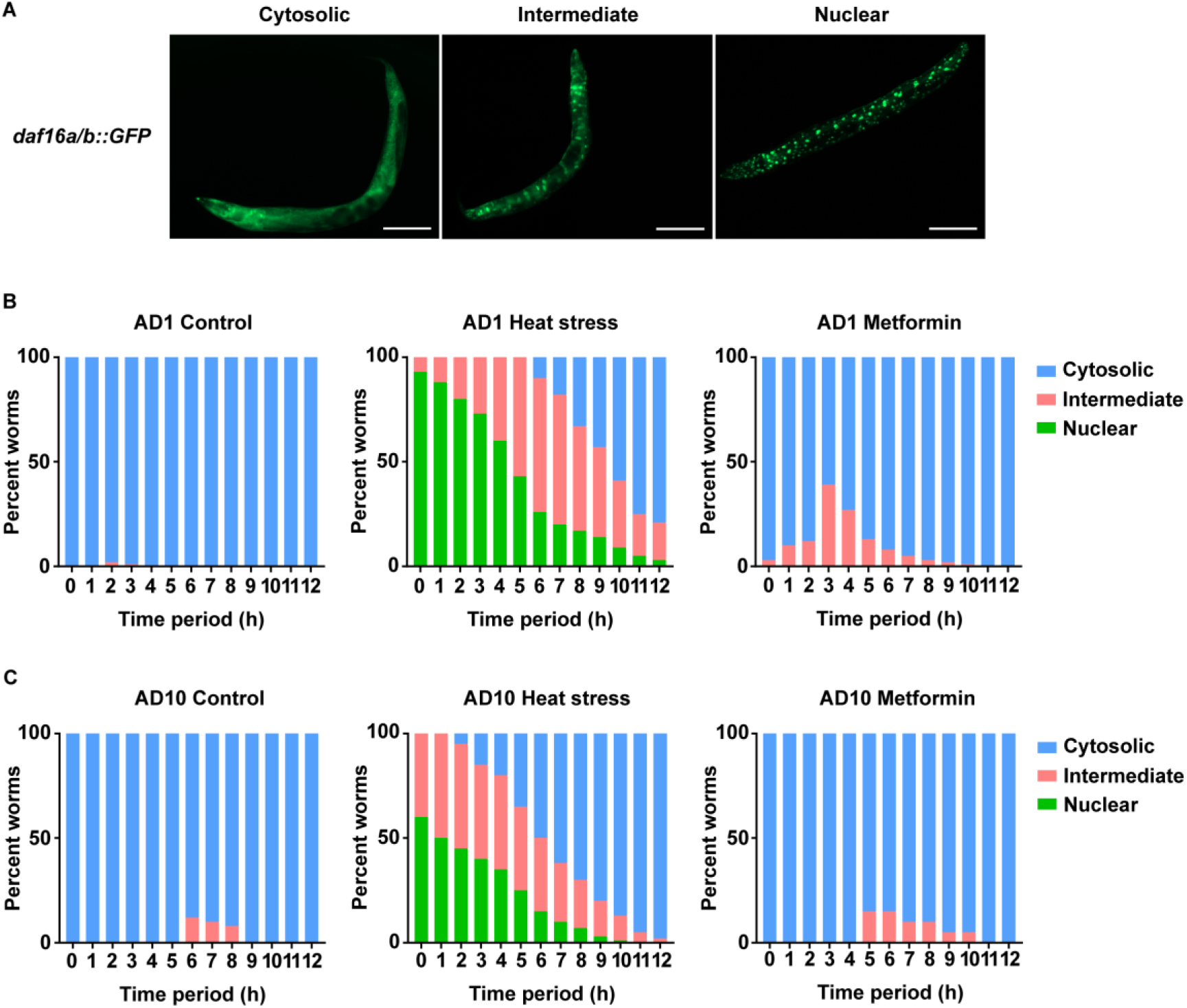
Metformin doesn’t induce DAF-16/FOXO nuclear translocation. Young (AD1) (**B**) and old (AD10) (**C**) transgenic nematodes expressing DAF-16::GFP fusion protein were treated with 50mM metformin or transient heat stress and DAF-16 sub-cellular localization was assessed by microscopy every hour; % of worms with specific localizations were quantified, n=100 for each condition, 4 independent tests were done and one representative result is shown. (**A**) representative images of DAF-16::GFP sub-cellular localizations used for the assessment in **B** and **C** are shown, scale bar is 100µm.

**Figure S9.**
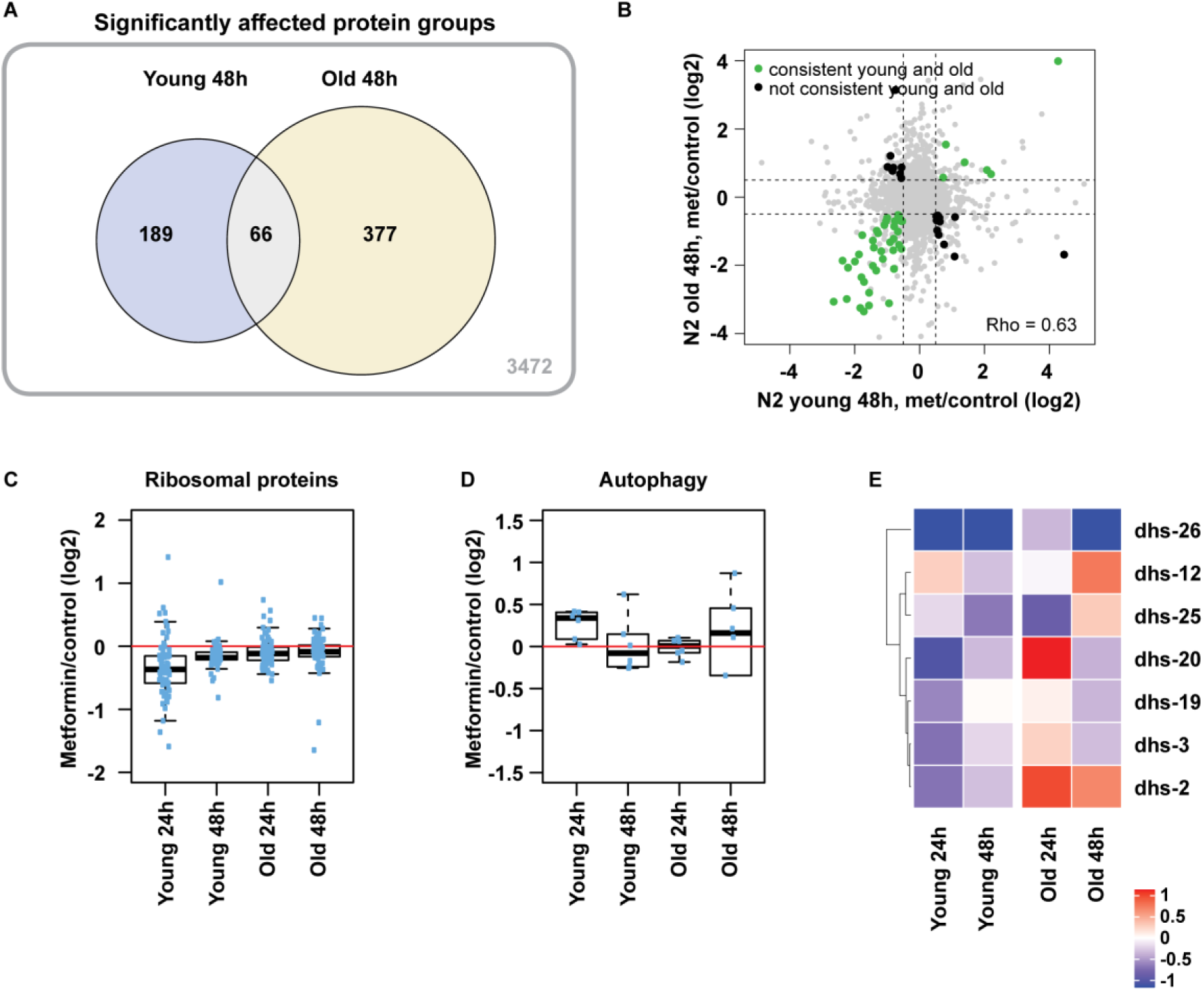
The induction of longevity assurance pathways and metabolic adaptations by metformin is age-dependent. Wild type worms were treated with 50mM metformin on adulthood day 1 (young) and 10 (old). (**A**) Venn diagram showing overlap between significantly altered proteins after 48h of early and late life metformin treatment is shown; the circled values include all proteins with absolute log2 fold change above 0.5 and Q value below 0.25; the number in the outside box indicates non-regulated proteins detected in both young and old samples collected at 48h; fold changes were calculated against age- and time point matched untreated controls. Three independent pools of worms were analyzed for each sample group. (**B**) Scatter plot comparing log2 fold changes of individual proteins (shown as dots) after 48h of metformin treatment at young and old age is shown. Proteins significantly regulated at both ages are highlighted with color: the green-highlighted proteins are consistently regulated at both ages while black-highlighted ones are regulated in the opposite direction between young and old age. The correlation coefficient (Rho) between young and old responses taking into account all regulated proteins is shown. Boxplots showing fold changes for selected ribosomal proteins (**C**) and proteins involved in general autophagy (**D**) are presented. The median fold change of the proteins belonging to each group is shown as a bold line, the upper and lower limits of the boxplot indicate the first and third quartile, respectively, and whiskers extend 1.5 times the interquartile range from the limits of the box. (**H**) Heatmaps of selected dehydrogenases are shown; only fold changes with significance in at least one age/treatment combination are depicted. n≥500 per replica and 3 replicas were measured for each condition. The list and complete data of proteins used for each boxplot and the heatmap are reported in Table S3.

**Figure S10.**
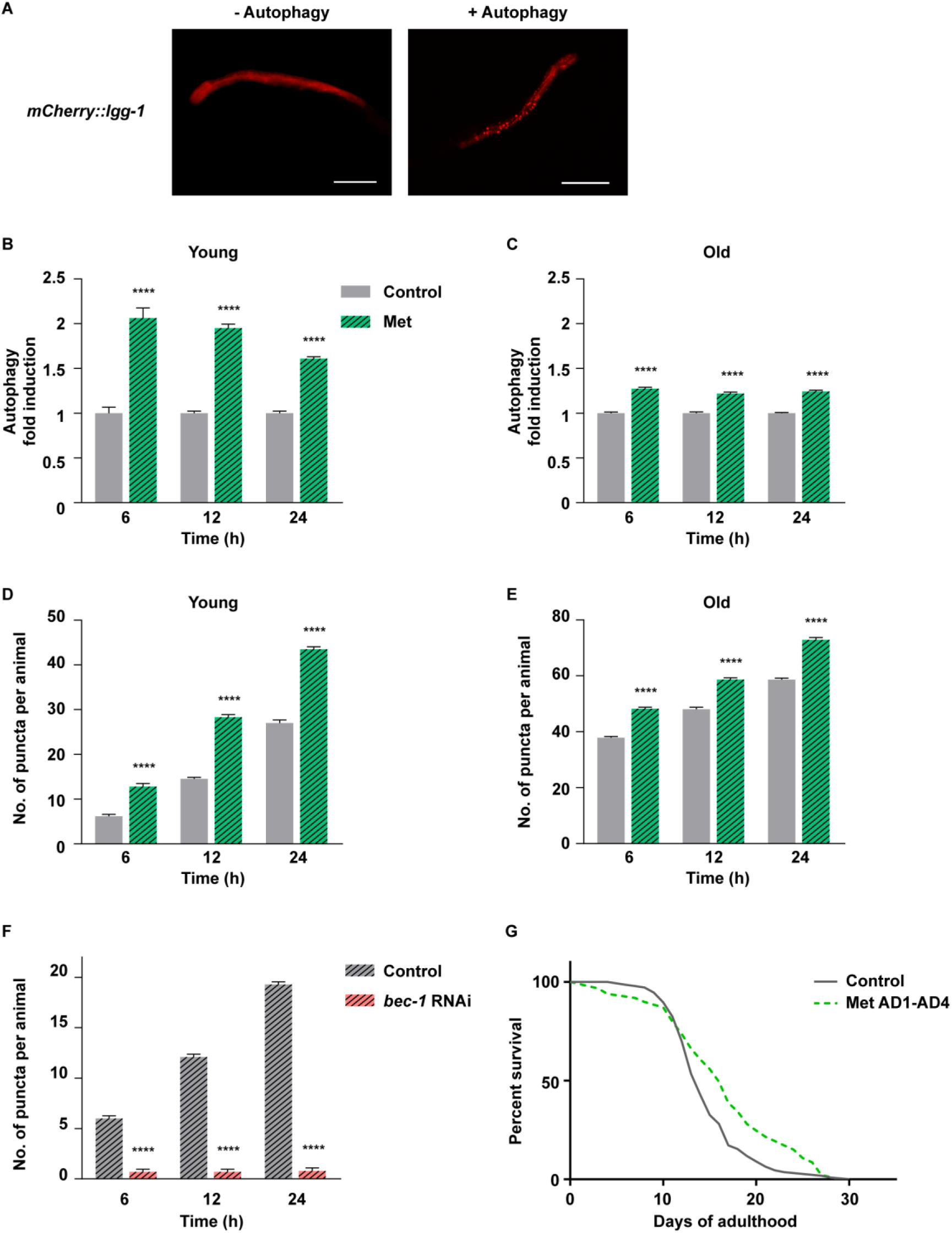
Induction of autophagy by metformin is impaired in late life. (**A**) Representative images of baseline diffused mCherry::LGG-1 expression **(left panel)** and autophagy puncta **(right panel)** are shown, scale bar is 100µm. Young (adulthood day 1, AD1) and old (AD10) transgenic animals were treated with 50mM metformin (Met), the number of puncta per animal was quantified, **B** and **C** show autophagy fold induction relative to time point matched untreated control in each case; **D** and **E** show absolute numbers of puncta per animal used for the calculation of values shown in B-C. The baseline elevation of autophagy over time is likely due to fresh plate transfer in all cases. (**F**) Transgenic animals were exposed to *bec-1* or control RNAi from the L4 larval stage; number of puncta was quantified over time. (**G**) Wild type nematodes were treated with 50mM metformin from day 1 (AD1) till day 4 (AD4) of adulthood, survival was scored daily. For **G** n≥100, significance was measured by log-rank test, the exact n numbers and statistical values are presented in Table S1. For **B-F** n=10, mean and SEM are presented, two-tailed unpaired t-test was used for the statistical analysis, all statistical values are presented in Table S2; **** p<0,0001.

**Figure S11.**
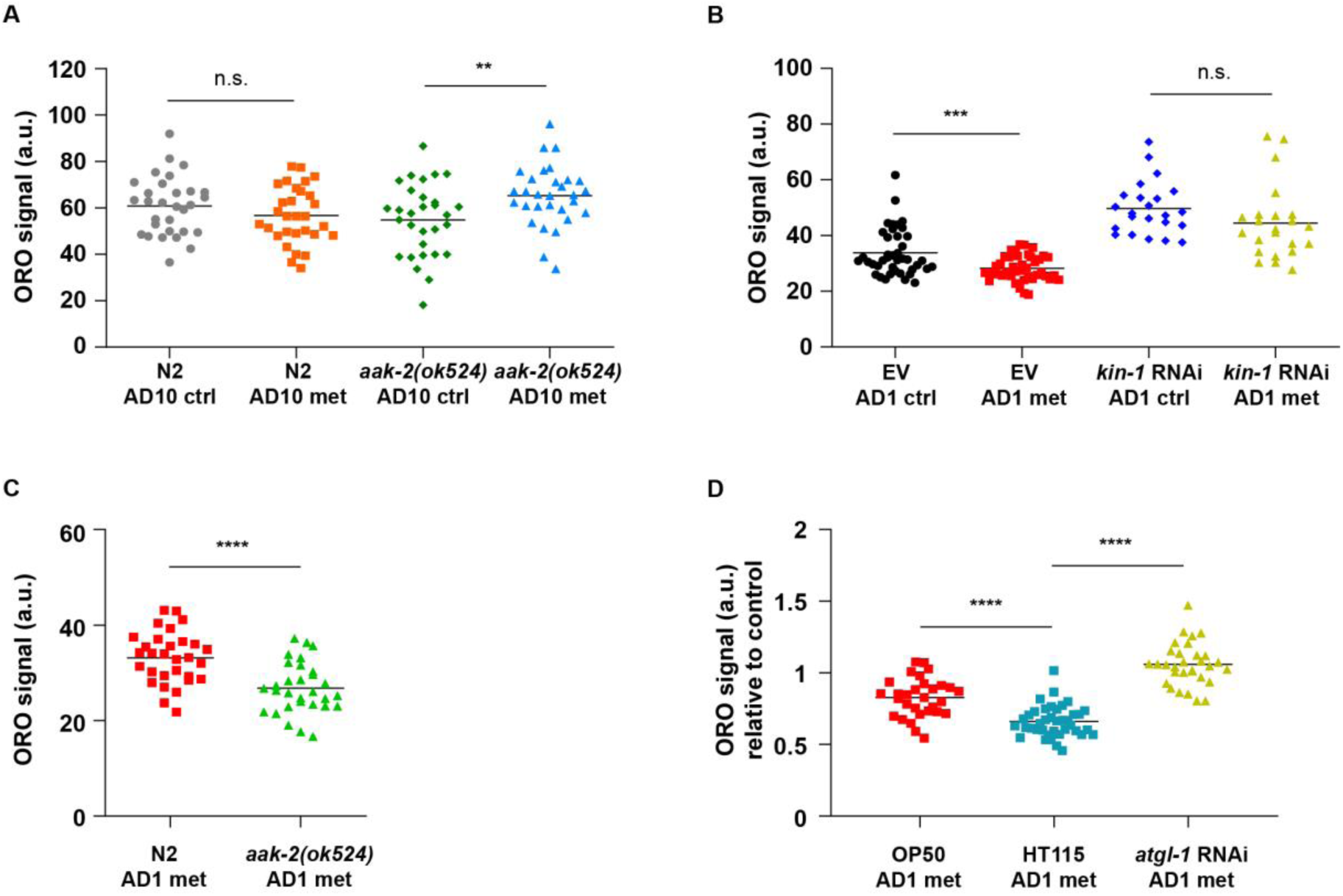
The age-specific effects of metformin on lipid reserves are regulated by PKA and AMPK pathways. (**A**) Wild type (N2, Bristol strain) and AMPK deficient *aak-2(ok524)* animals were treated with 50mM metformin on adulthood day 10 (AD10), whole body Oil Red O lipid staining was performed after 24h of treatment. (**B**) Pre-adult (larval stage 4) wild type animals were exposed to empty vector RNAi (EV) or *kin-1* RNAi for 12h prior to treatment with 50mM metformin. The limited RNAi exposure was necessary because early age *kin-1* RNAi treatment had a strong effect on nematode development. Oil red O whole body lipid staining was performed after 24h of metformin exposure. (**C**) Wild type and AMPK deficient animals were treated with 50mM metformin on adulthood day 1 (AD1) and whole body Oil Red O lipid staining was performed after 24h of treatment. The absolute difference between metformin exposed mutant and control animals is presented (complementary to Fig 6E), demonstrating a stronger metformin-triggered decline of lipid content in AMPK deficient animals. (**D**) Young wild type animals were treated with 50mM metformin in the presence of OP50 *E. coli*, HT115 *E.coli* containing the empty vector and *atgl-1* RNAi provided in HT115 bacteria; whole body Oil Red O lipid staining was performed after 24h of treatment, presented values were normalized to respective metformin free control in each case. Oil Red O mean grey values were taken as arbitrary units (a.u.) in all cases. For **A** and **C** n=30, for **B** n≥23 and for **D** n≥30; in all cases mean value is presented, two-tailed unpaired t-test was used for the statistical analysis, all statistical values are presented in Table S2; ** p<0,01; *** p<0,001; **** p<0,0001.

**Figure S12.**
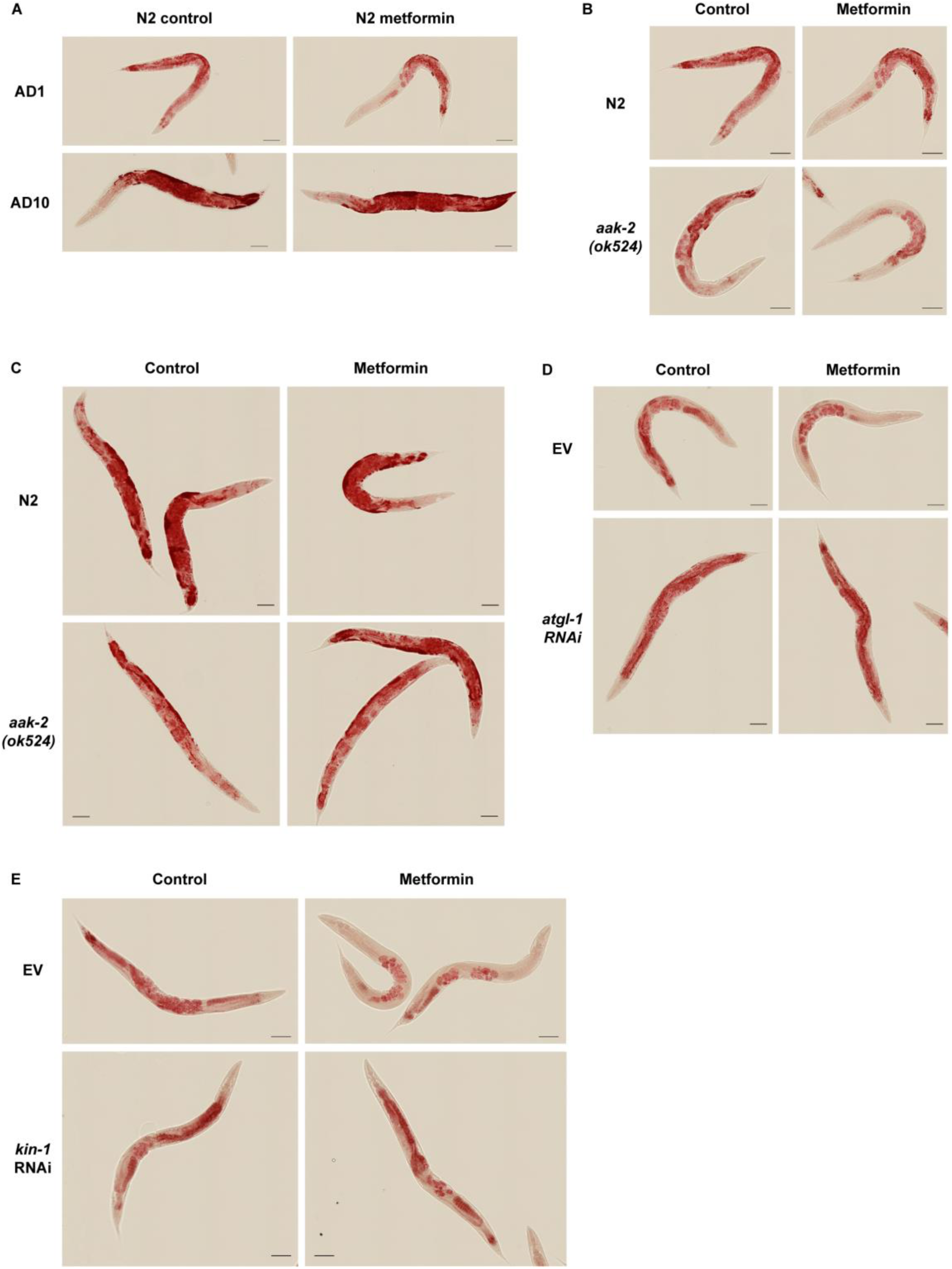
Lipid levels were assessed by whole body Oil Red O staining in nematodes. Representative images corresponding to Figures 6A (young and old wild type animals) **(A)**, 6E (young animals) **(B)**, S11A (old animals) **(C)**, 6F (young animals) **(D)** and S11B (young animals) **(E)** are shown. For (**A**-**E**) scale bar is 100µm. For **A**-**C** n=30, for **D** n≥31 and for **E** n≥23.

**Figure S13.**
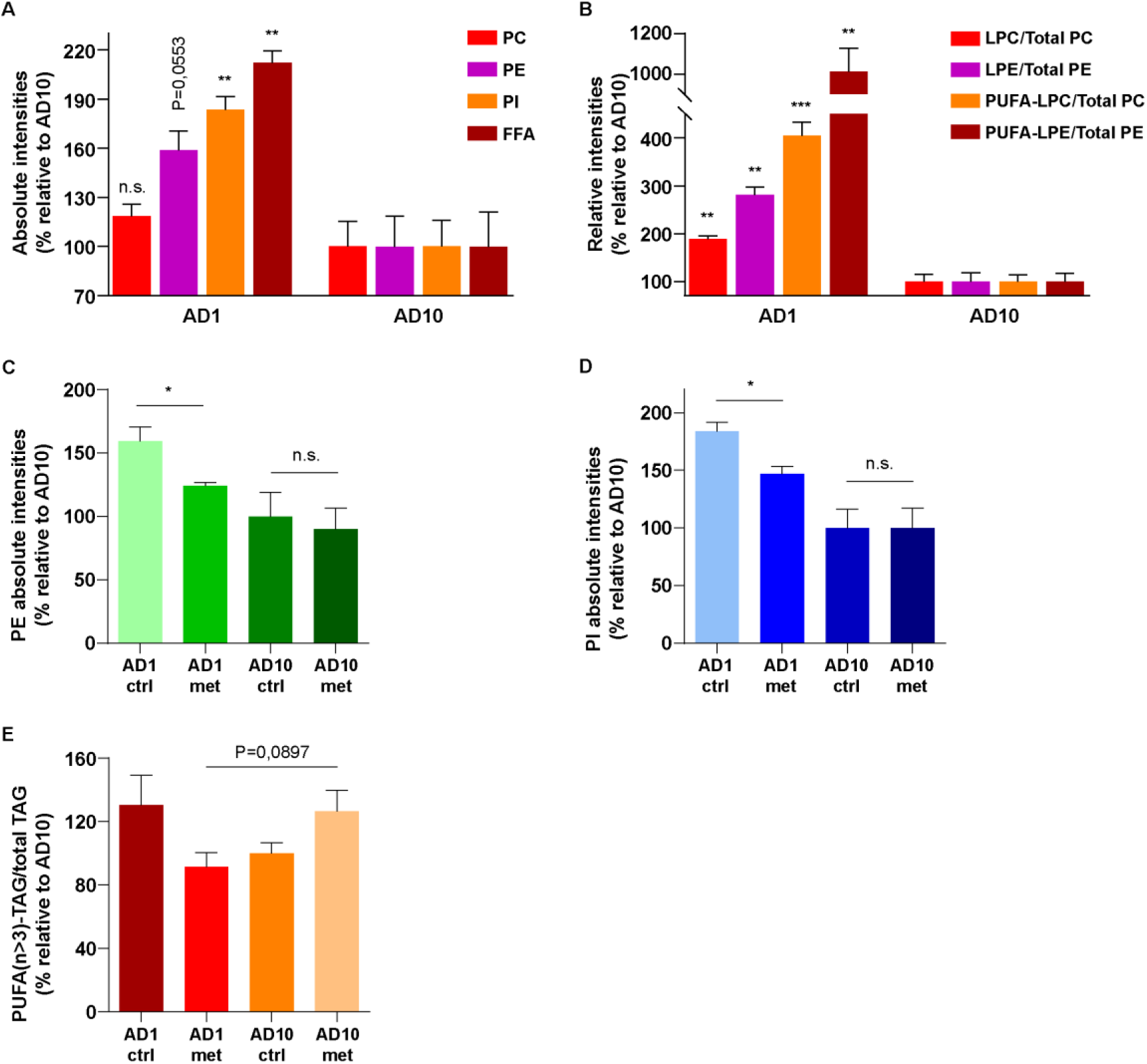
The dietary restriction mimetic lipid turnover response to metformin is blunted in late life. Wild type animals were treated with 50mM metformin for 24h on AD1 and AD10; lipids were isolated and analyzed by UPLC-MS/MS. Absolute intensities for phosphatidylethanolamines (PEs) (**C**) and phosphatidylinositols (PIs) (**D**) are shown for treated and untreated animals, and absolute intensities of PEs, PIs, free fatty acids (FFAs) and phosphatidylcholines (PC) are shown for untreated young and old animals (**A**). (**B**) Relative intensities of lyso-phosphatidylcholines (LPCs) and lyso-phosphatidylethanolamines (LPEs) as well as of polyunsaturated fatty acid (PUFA) containing LPCs and LPEs are shown for untreated young and old animals. (**E**) Relative intensities of triglycerides with PUFAs containing more than 3 double bonds (PUFA n>3 TAGs) are shown for young and old treated and untreated animals. All values are normalized to the AD10 untreated control. Definitions of absolute and relative intensities are provided in the Methods section. For **A-E** 700 animals were analyzed in 3 replicas for each condition, all individual lipid values are presented in Table S4, mean and SEM are depicted, two-tailed unpaired t-test was used for the statistical analysis, all statistical values are presented in Table S2; * p<0,05; ** p<0,01; *** p<0,001.

**Figure S14.**
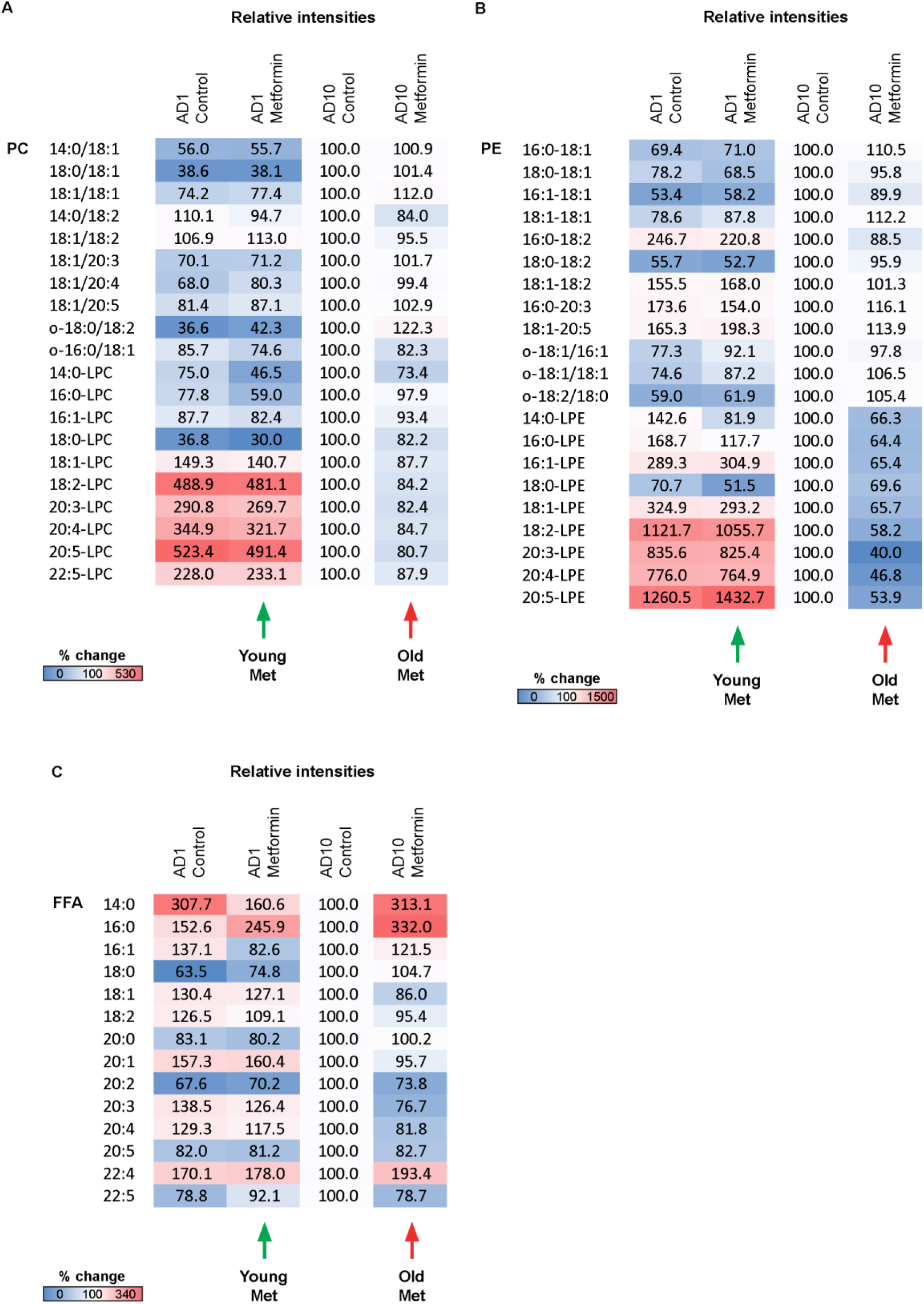
Metformin treatment triggers age specific patterns of lipid utilization. Wild type animals were treated with 50mM metformin for 24h on AD1 and AD10; lipids were isolated and analyzed by UPLC-MS/MS. Mean relative intensities of distinct phosphatidylcholines (PC) (**A**), phosphatidylethanolamines (PEs) (**B**), and free fatty acids (**C**) are presented. All values are normalized to the AD10 untreated control. For **A-C** 700 animals were analyzed in 3 replicas for each condition, all individual lipid values are presented in Table S4.

**Figure S15.**
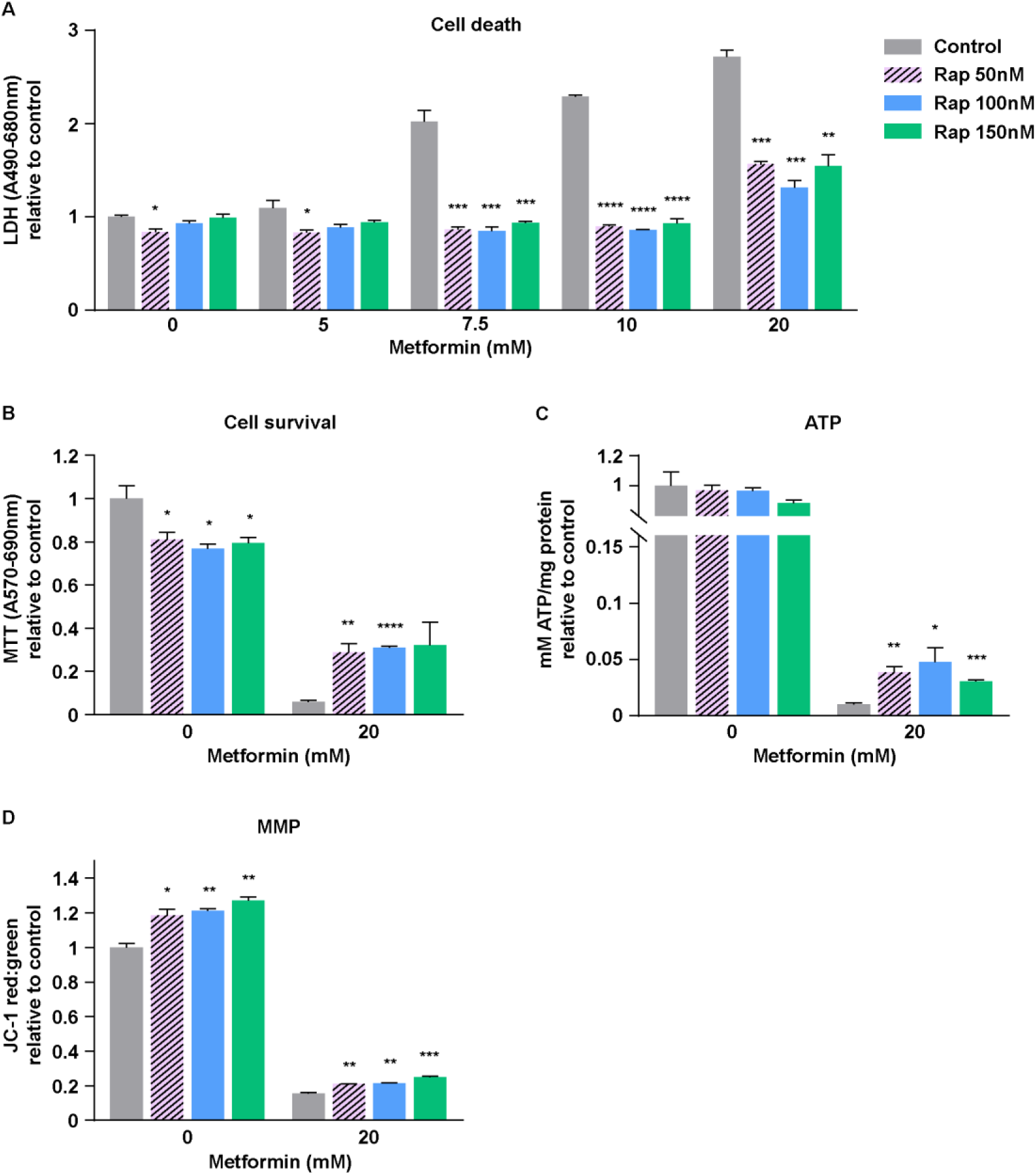
Rapamycin co-treatment alleviates metformin toxicity in human primary cells. Pre-senescent (PD44) primary human skin fibroblasts were treated with indicated doses of metformin in presence or absence of indicated concentrations of rapamycin (Rap) for 24 (**A, B**) and 20 (**C, D**) hours. Cell death (**A**, LDH assay), cell survival (**B**, MTT assay), ATP content (**C**) and mitochondrial membrane potential (**D**, JC-1 staining) were measured; the data are complementary to Fig. 7A-C; values are relative to the respective untreated control (no rapamycin, no metformin) for each assay. n=3, mean and SEM are presented, two-tailed unpaired t-test was used for the statistical analysis, statistical values are shown in Table S2; * p<0,05; ** p<0,01; *** p<0,001; **** p<0,0001.

**Figure S16.**
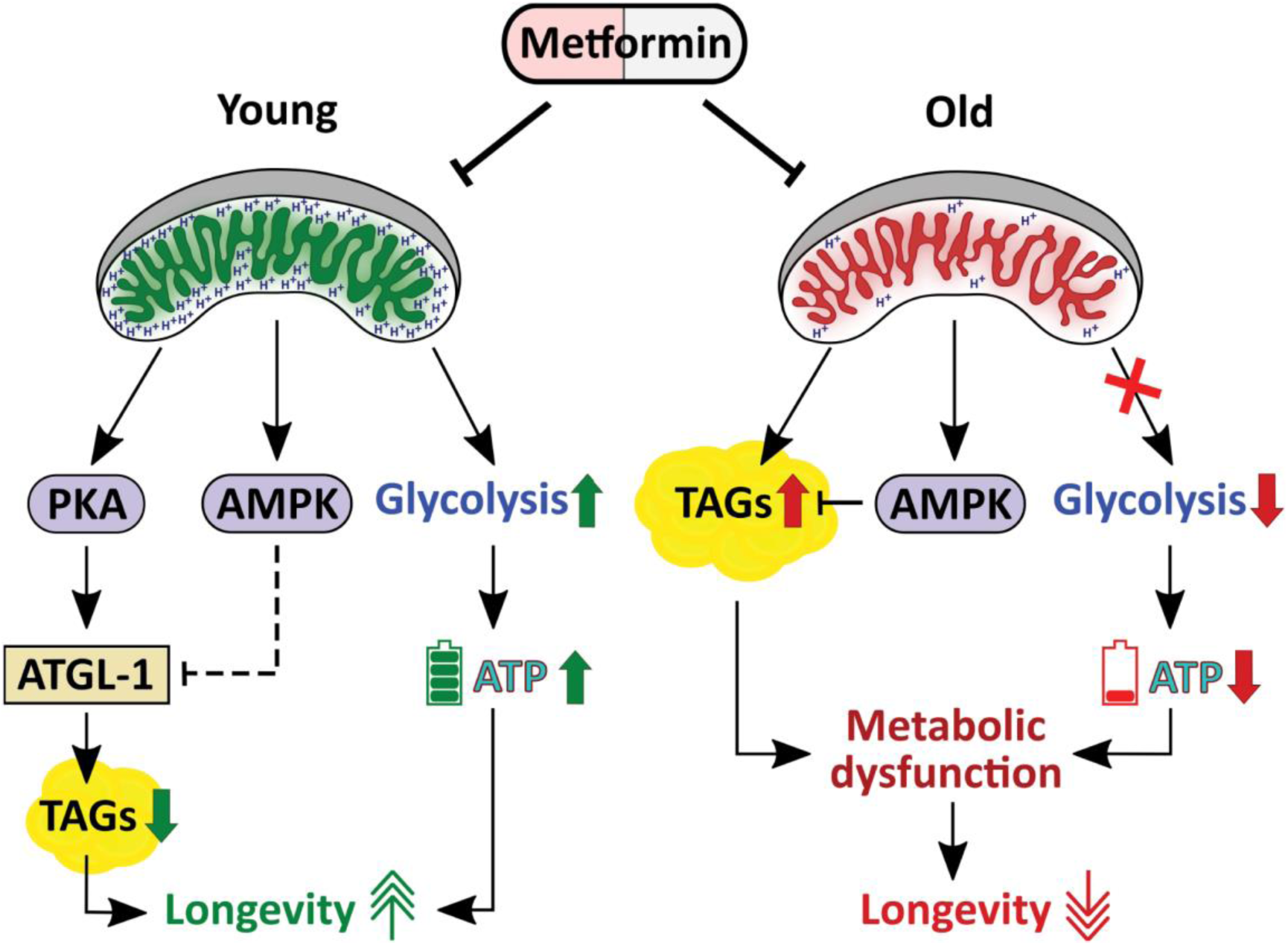
Metformin exerts age-specific effects on energy metabolism and instigates a severe energetics failure in late life. While early life metformin treatment triggers metabolic adaptations, such as elevated glycolysis and the DR-like utilization of lipids, which support the longevity benefits of this drug, late life metformin exposure acts in concert with aging-associated metabolic distortions to trigger a sever metabolic failure culminating in ATP exhaustion incompatible with cell viability. Importantly, the early life DR effect of metformin is executed by the PKA pathway while AMPK likely prevents the untimely lipid expenditure by this pathway to ensure long-term longevity benefits of metformin.

**Table S1.**
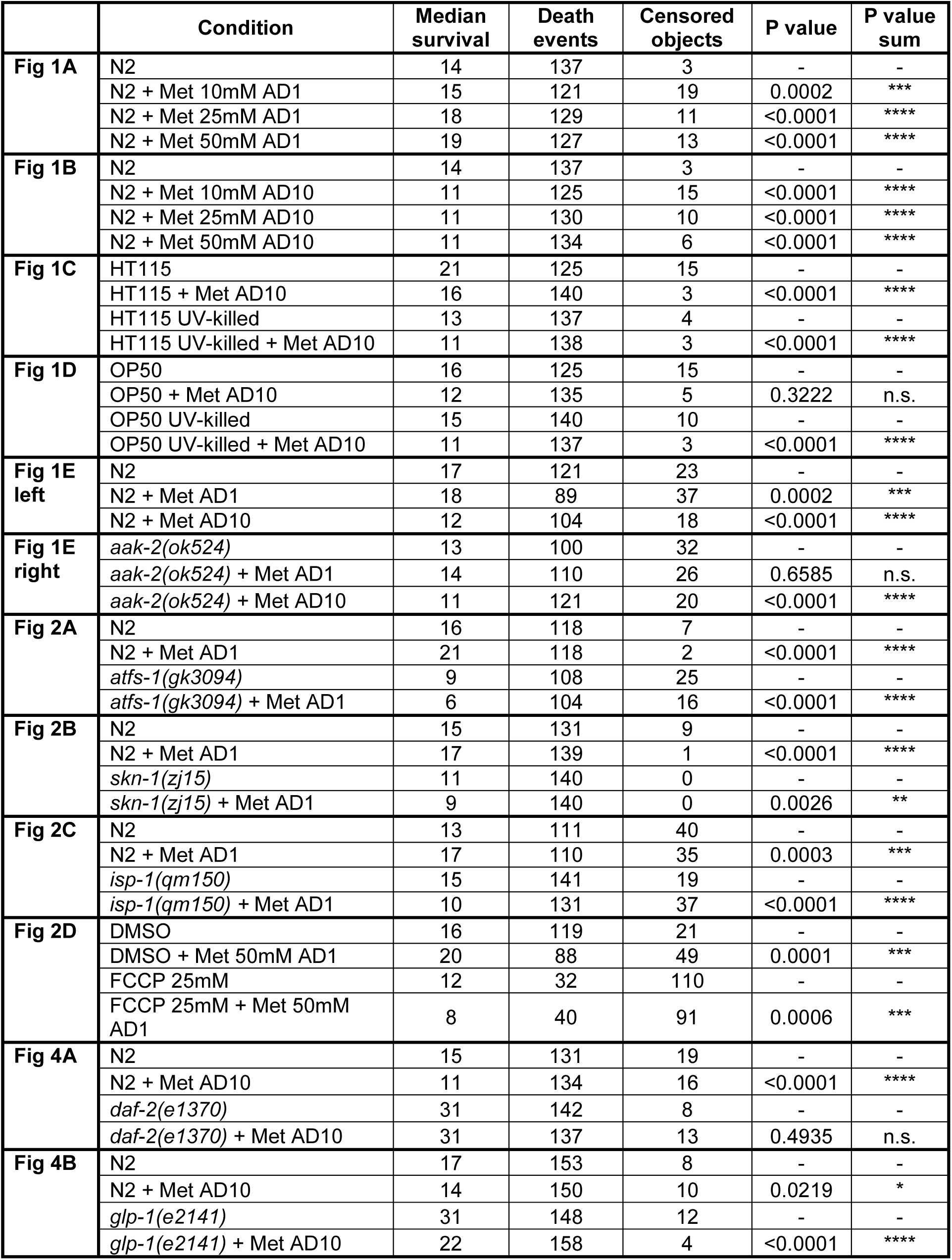
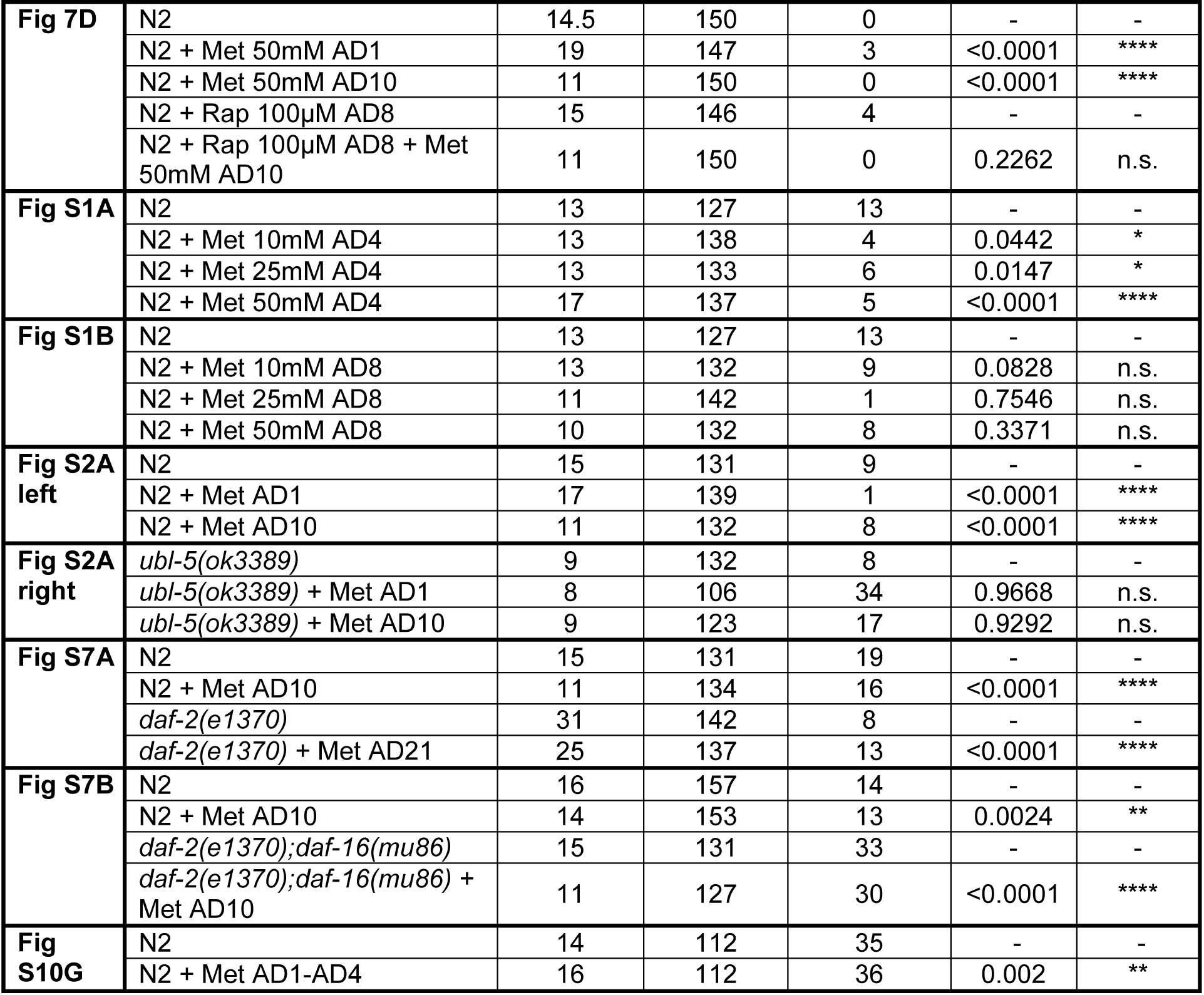
Statistics for lifespan experiments (log rank test).

**Table S2.**
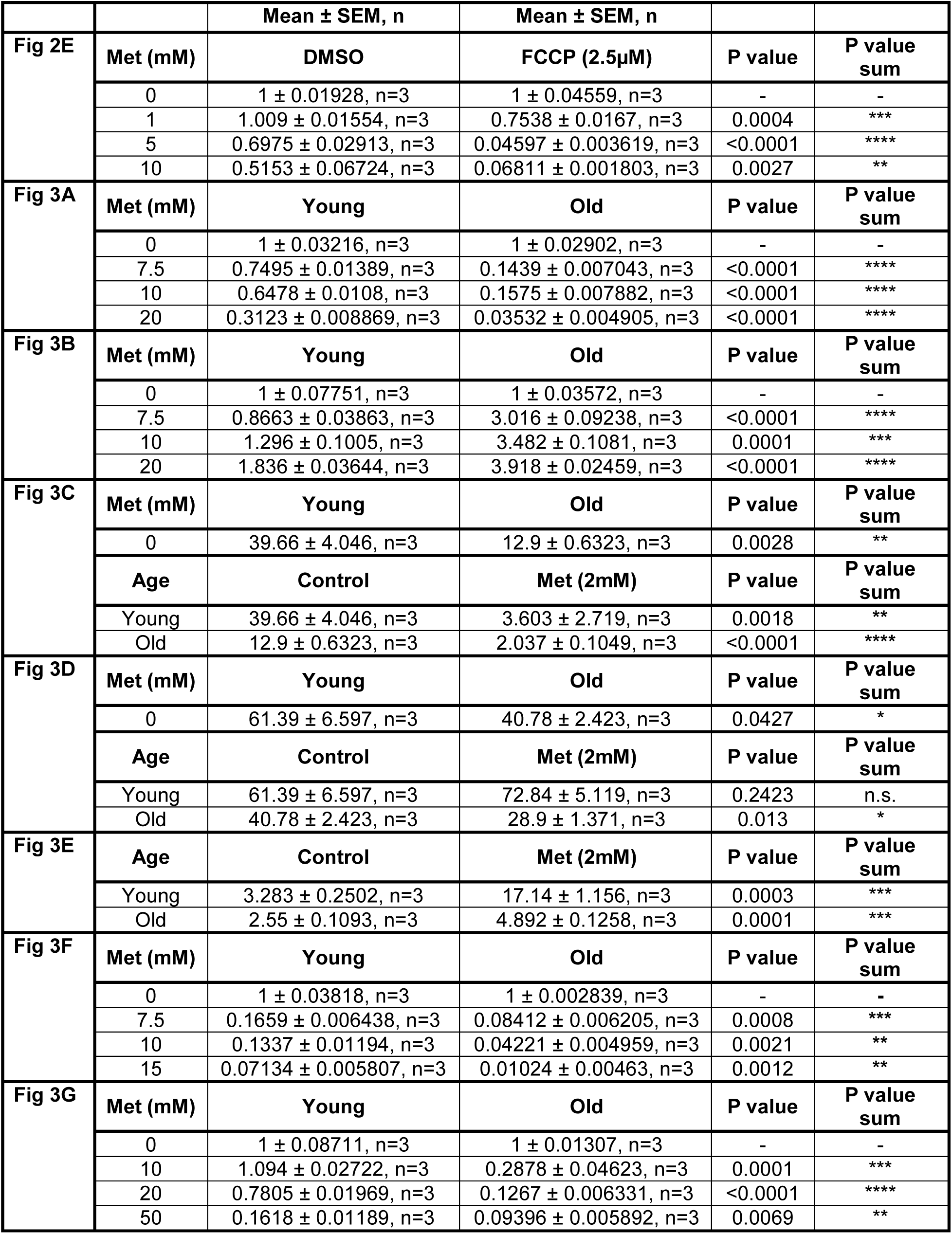
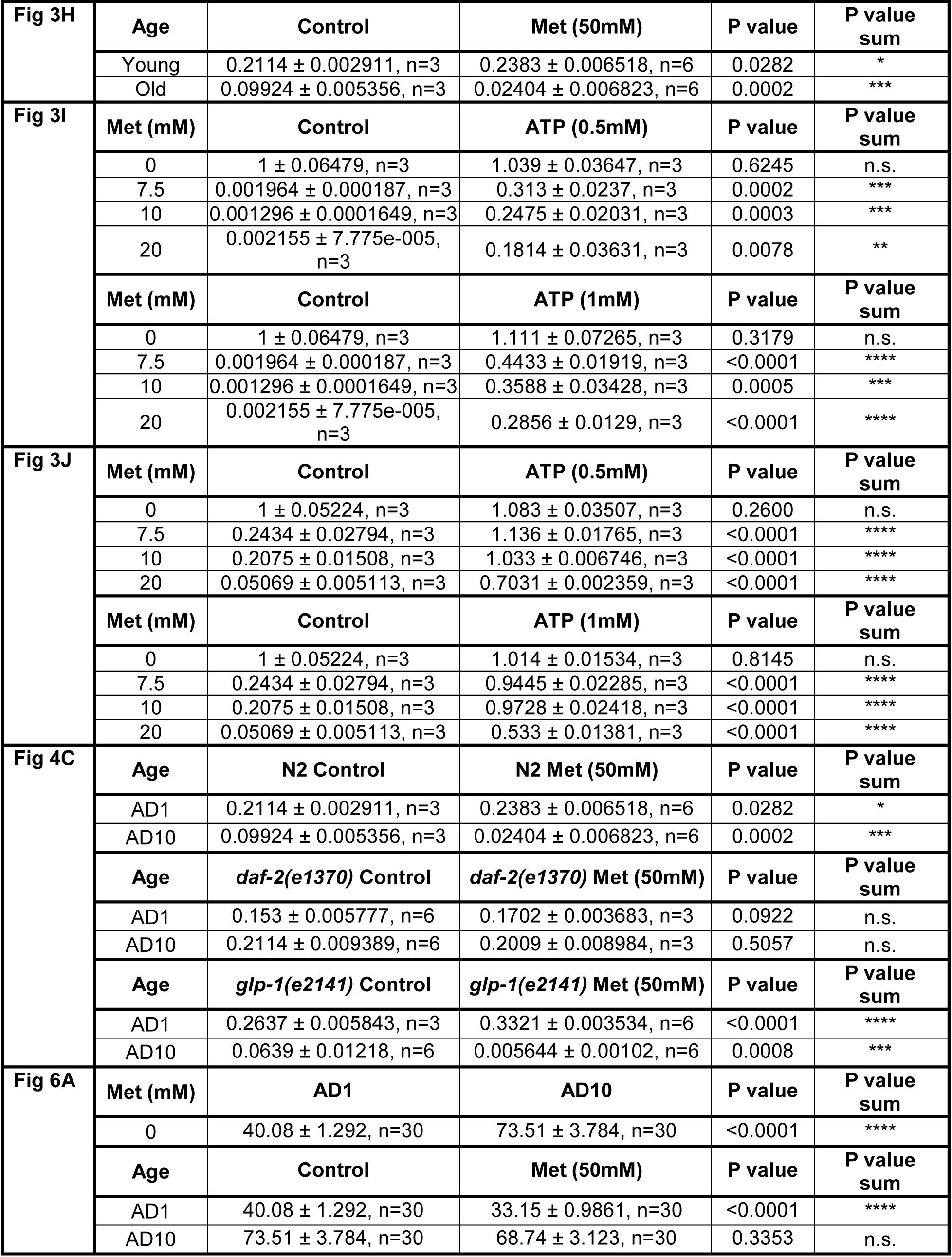
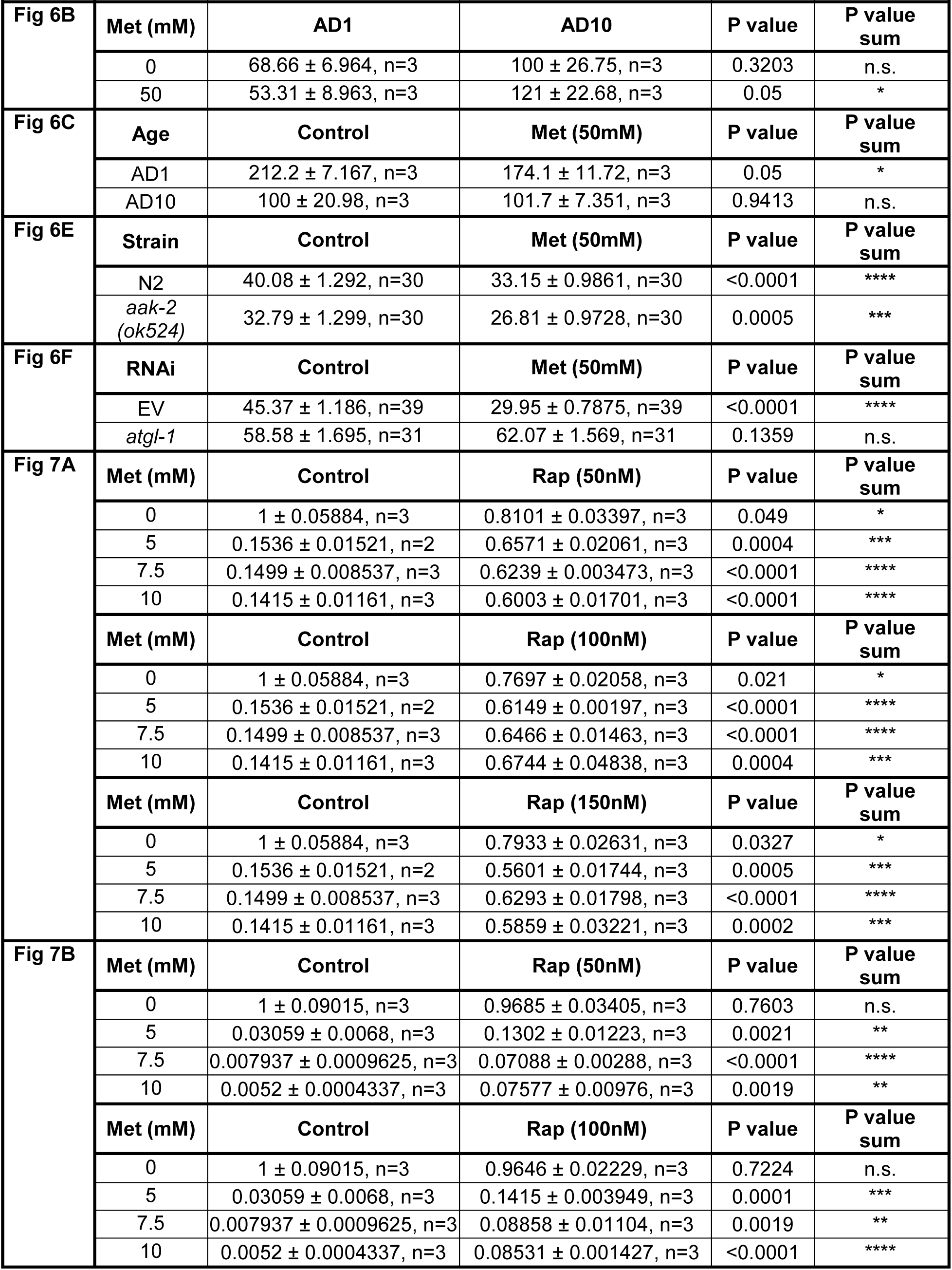
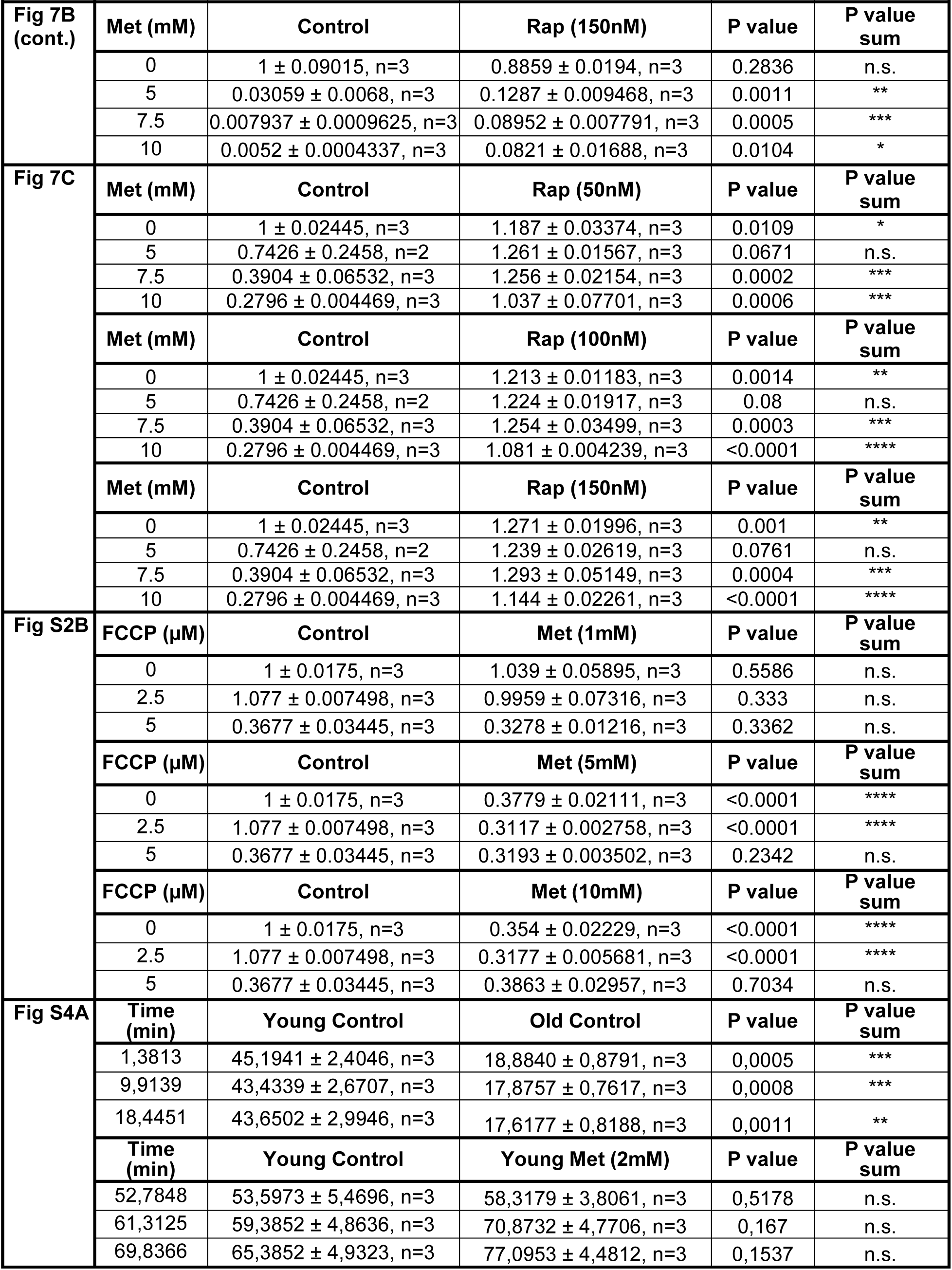
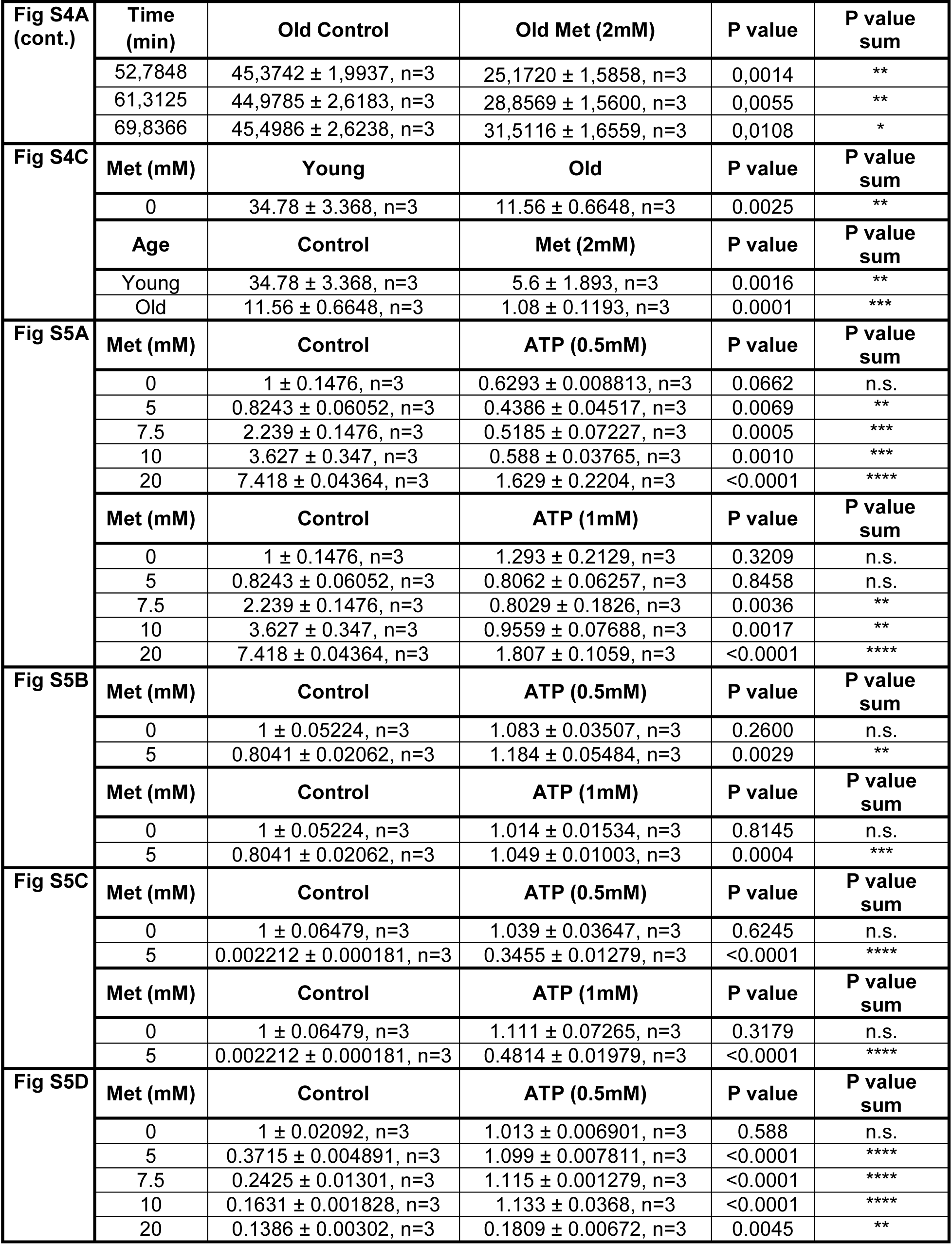
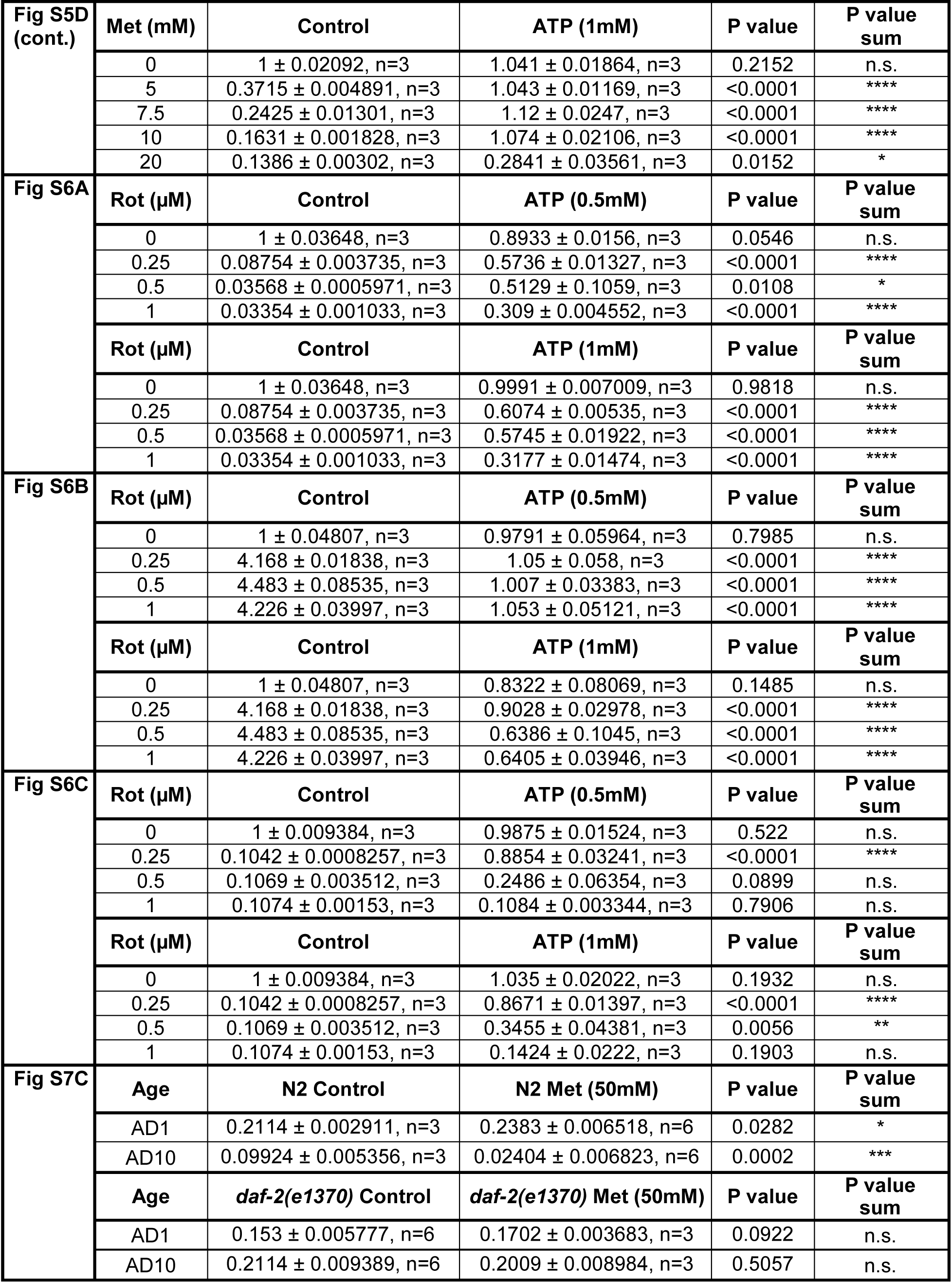
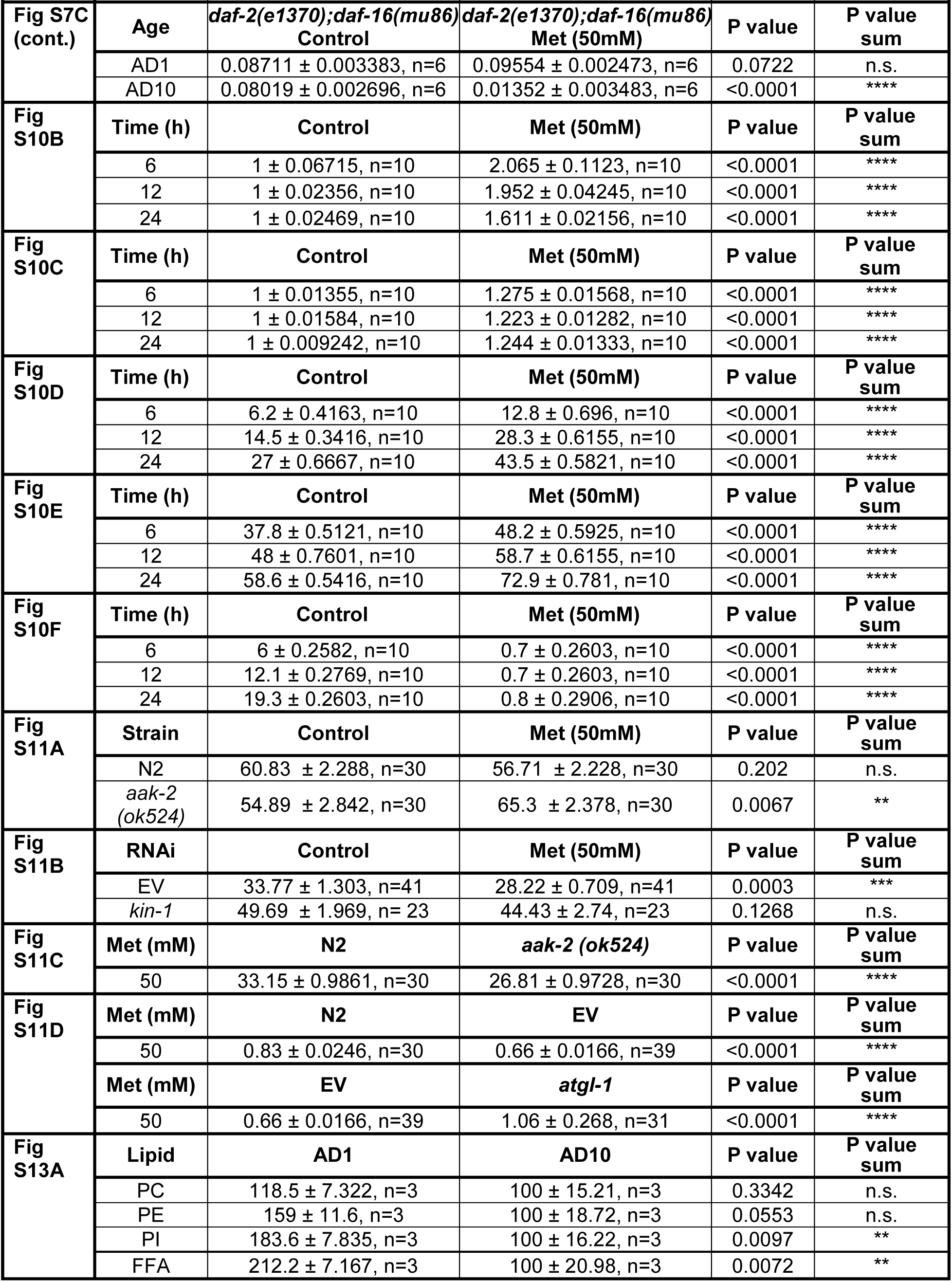
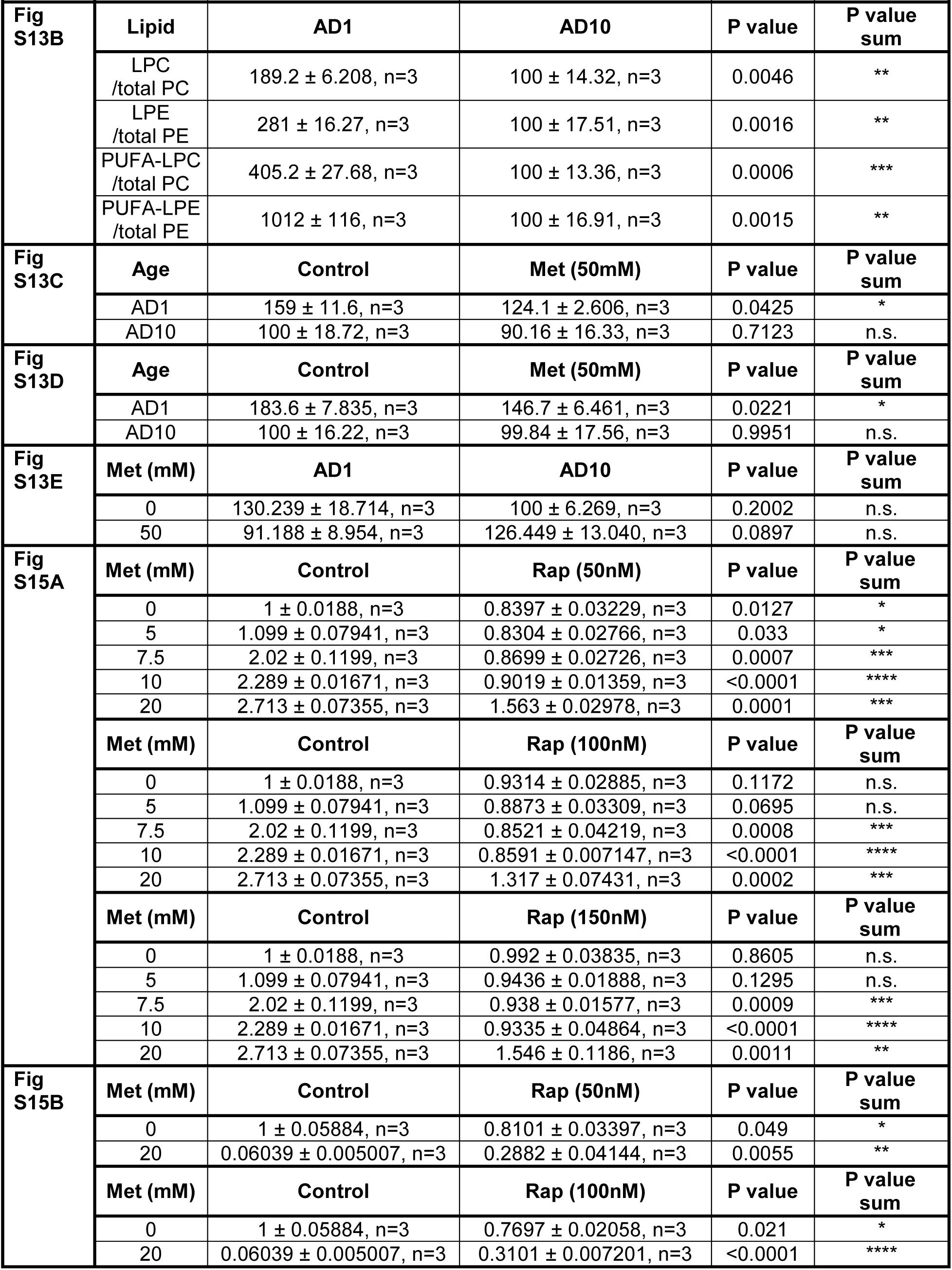
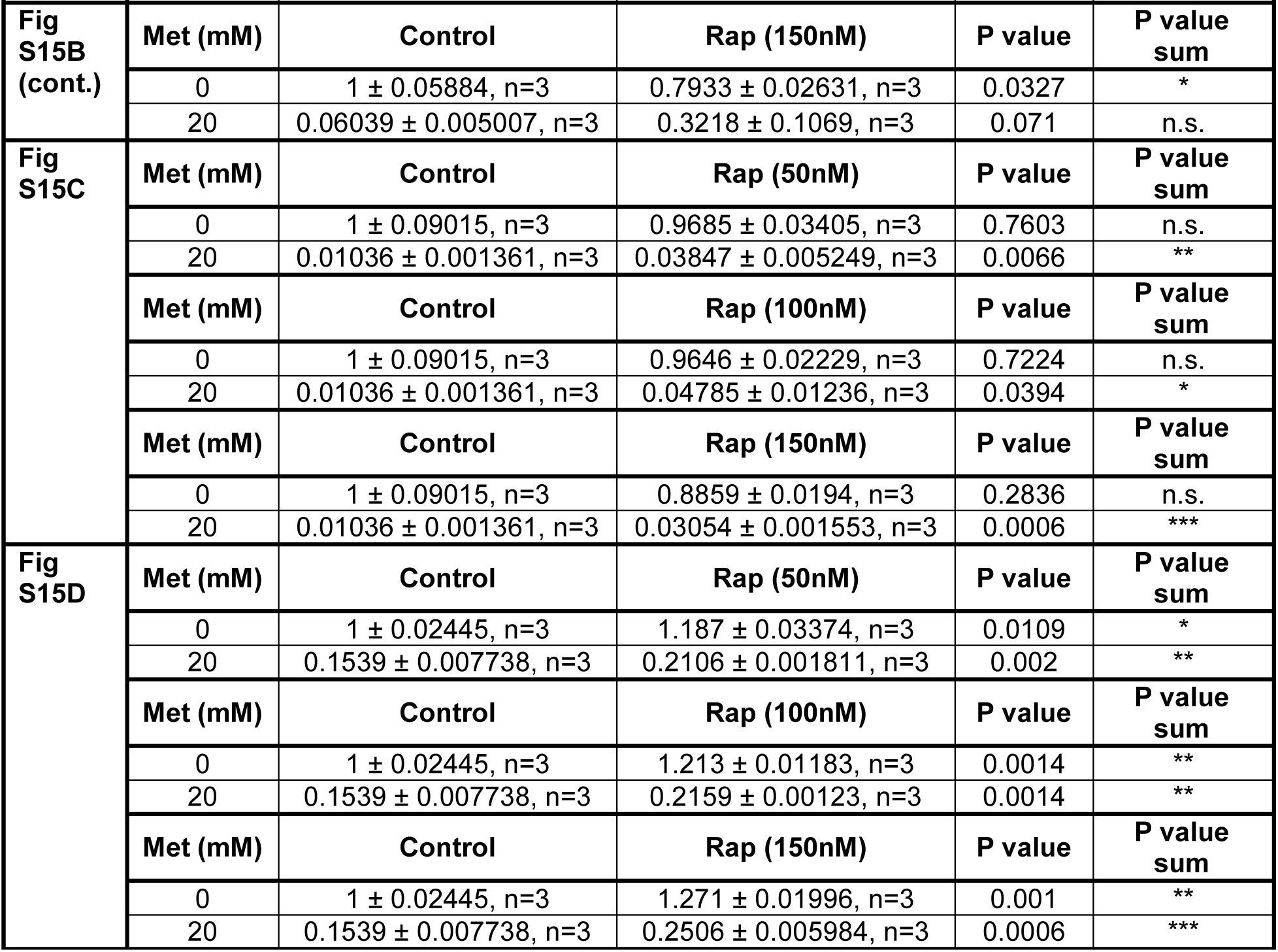
Statistics for all data analyzed by t-test.

**Table S3. Proteomics data analysis (separate file).**

The first tab of the table shows individual fold changes of all proteins measured in metformin treated samples relative to the age and time point matched untreated control; statistical values are included in each case and the fold changes are color-coded. Individual tabs summarize protein identities and statistical values used to generate the corresponding figures; the last (to the right) column of the tab depicts the assignment of proteins to distinct pathway groups. Tab 4 depicts an independent experiment performed in order to assess the mitochondrial content in respective genetic backgrounds.

**Table S4. Lipidomics data analysis (separate file).**

The table lists all raw intensities used and explains the calculations which connect these values to presented figures.

